# The production of diverse brGDGTs by an Acidobacterium allows a direct test of temperature and pH controls on their distribution

**DOI:** 10.1101/2022.04.07.487437

**Authors:** Yufei Chen, Fengfeng Zheng, Huan Yang, Wei Yang, Ruijie Wu, Xinyu Liu, Huayang Liang, Huahui Chen, Hongye Pei, Chuanlun Zhang, Richard D. Pancost, Zhirui Zeng

## Abstract

Microbial lipid biomarkers preserved in geological archives can be used to explore past climate changes. Branched glycerol dialkyl glycerol tetraethers (brGDGTs) are unique bacterial biomarkers that have been used as molecular tools for the quantitative determination of terrestrial temperatures and the pH of depositional environments over a range of geological timescales. However, the exact biological source organisms – especially of the entire suite of brGDGTs found in the environment – remains unclear; by extension, so do the mechanisms that govern these proxies. Here, we identified a brGDGT-producing strain *Candidatus Solibacter usitatus* Ellin6076, by identifying archaeal tetraether synthase homologs in bacterial genomes. This strain synthesizes diverse brGDGTs, including regular C_5_-methylated and cyclic brGDGTs, and brGDGTs comprise up to 66% of the major lipids, far exceeding the proportions found in previous studies. The degree of C_5_-methylation in cultured strain Ellin6076 is primarily determined by temperature, whereas cyclization appears to be influenced by multiple factors. Consequently, culture-derived paleoclimate indices are in agreement with the global soil-derived MBT’_5ME_ (methylation index of C_5_-methyl brGDGTs) proxy for temperature but not the CBT_5ME_ (cyclization index of C_5_-methyl brGDGTs) proxy for pH. Our findings provide important insights from a physiological perspective into the underlying mechanism of brGDGT-based proxies.

**Significance Statement:** Branched glycerol dialkyl glycerol tetraethers (brGDGTs) are biomarkers widely used for the quantitative estimation of past climatic changes due to their ubiquitous occurrence in the environment and the relationships between their distributions and temperature and pH. However, the ecophysiology of brGDGT-producing bacteria and the mechanistic basis for brGDGT-based climate proxies remain unknown. Here, we identify a brGDGT-producing Acidobacterium and present a physiological study of brGDGTs in response to cultivation variables, which provides pivotal insights into how brGDGT producers modulate methylation and cyclization under different culturing conditions. Our study represents a significant advancement in understanding the physiological role of lipid structures in microbial adaptation and helps us interpret the relationships between brGDGT-based proxies and environmental conditions of the geological environment.

## Introduction

Quantitative estimation of past climate change is important for understanding Earth history, contextualising the impact of recent human-induced climate change, and testing models used for future projections. This is challenging, particularly for the terrestrial environment, due to the scarcity of quantitative proxies for the reconstruction of climate variables, e.g. temperature and precipitation. Microbial lipid biomarkers preserved in terrestrial climate archives offer several useful tools for documenting the evolution of Earth’s climate (1). Branched glycerol dialkyl glycerol tetraethers (brGDGTs) are one class of such lipids and have been used to reconstruct past temperature, paleohydrology, pH, and terrigenous organic input (2–5). Due to their ubiquitous occurrence in terrestrial and aquatic settings, the number of their applications to climate archives, e.g. paleosols, peats, lake sediments, stalagmites, estuarine and marine sediments, has increased dramatically over the last decade (3, 6).

These applications rest on the empirical relationships between the distribution of brGDGTs and environmental variables, such as temperature, mean annual precipitation, and pH in modern soils and surface sediments (7–9). In particular, considerable efforts have been devoted to improving our understanding and the accuracy of brGDGT-based temperature or pH proxies (2, 10, 11). However, uncertainties persist, many arising from the fact that the microbial producers of these mysterious lipids in the environment remain incomplete (12, 13). This has prevented an examination of the ecophysiology of these microbes and testing of these proxies under laboratory conditions.

The quest for the microbial producer(s) of brGDGTs has been ongoing since the discovery of brGDGTs almost twenty years ago (14). The enantiomeric configuration of the glycerol backbone, 1,2*-*di*-O-*alkyl*-sn-*glycerol, assigns brGDGTs as lipids that are synthesized by bacteria (15). A paired 16S rRNA gene sequencing and brGDGT approach was used in numerous studies to help constrain the identity of these bacteria (16–18), yielding a range of bacterial phyla, such as Acidobacteria, Bacteroidetes, and Verrucomicrobia, as potential producers of brGDGTs in the environment (16–18). Among these microbes, Acidobacteria were suspected to be the most likely biological sources of brGDGTs (12), because particularly high abundances of brGDGTs correspond with the dominance of Acidobacteria in the bacterial community in soils and peats (18). This was finally confirmed by the examination of the lipid profiles of more than 40 Acidobacterial strains, revealing their widespread production of the potential building block (*iso*-diabolic acid) for brGDGTs, and, most importantly, the positive identification of brGDGT-Ia, a tetramethylated brGDGT, in *Edaphobacter aggregans* Wbg-1 and *Acidobacteriaceae* bacteria A2-4c (12, 19, 20). Subsequent work has shown that oxygen limitation can trigger the production of more brGDGT-Ia in *E. aggregans* (13), likely explaining the long-observed association of brGDGTs with low oxygen conditions. However, the majority of brGDGTs that are used in climate proxies were absent from the lipid profiles of Acidobacteria previously examined in cultures (12, 19, 20). This necessitates a further search for the biological source(s) of brGDGTs in the environment.

Recently, tetraether synthase (Tes), a key protein responsible for the formation of archaeal isoprenoid GDGTs (isoGDGTs) via the combination of two archaeol molecules, has been identified (21). Bacterial brGDGTs bear a structural resemblance to archaeal isoGDGTs, i.e., both of them consisting of two alkyl chains linked to two glycerol backbones via four ether bonds (15). The Tes protein could, therefore, also be involved in the biosynthesis of bacterial brGDGTs, since Tes homologs have been found in bacterial genomes of diverse phyla including Acidobacteria (21).

In this study, we identified a brGDGT-producing strain *Candidatus Solibacter usitatus* Ellin6076, a member of Acidobacteria subdivision 3, through searching archaeal Tes homologs in bacterial genomes. Crucially, strain Ellin6076 can synthesize regular brGDGTs with more than 4 methyl groups and cyclopentane moieties as its major membrane lipids. This allows us to assess the physiological basis for the brGDGT responses to changes in temperature, pH, and oxygen level, which provides insights into the underlying mechanism of brGDGT-based paleoclimate proxies.

## Results

### Identification of brGDGTs in Acidobacteria culture

Following our hypothesis that the Tes homolog protein is associated with bacterial brGDGT production, we searched for Tes homologs in bacterial genomes to determine the potential biological source of brGDGTs. Strain Ellin6076 was noteworthy, as it has one Tes homolog with high sequence alignment scores (identity = 40%, e-value = 1e^-139^) with archaeal functional Tes (MA_1486). Moreover, it also contains a possible archaeal GDGT ring synthase (Grs) (22) homolog (identity = 25%, e-value = 1e^-40^). We cultured strain Ellin6076 aerobically under optimal growth conditions at 25 °C and pH 5.5 for 14 days, and then identified its lipid profile with reversed-phase–liquid chromatography–high-resolution mass spectrometer (RP–LC–HRMS) and normal-phase–liquid chromatography–mass spectrometer (NP–LC–MS). The results showed that strain Ellin6076 produced a series of brGDGT compounds, including brGDGT-Ia, Ib, Ic, IIa, IIb, IIc, and IIIa. The brGDGT-IIIb and IIIc components were not detected, perhaps due to their absence or concentrations below the detection limit. Compounds anticipated to be related to brGDGT biosyntheses, such as *iso*C_15_-dialkyl glycerol ether (DGE), branched glycerol trialkyl glycerol tetraethers (brGTGTs), and branched glycerol dialkanol diethers (brGDDs), were also detected. The intact polar lipids (IPLs) corresponding to some of the above core lipids were also identified (Fig. 1; *SI Appendix*, Fig. S1−3).

**Fig. 1.**
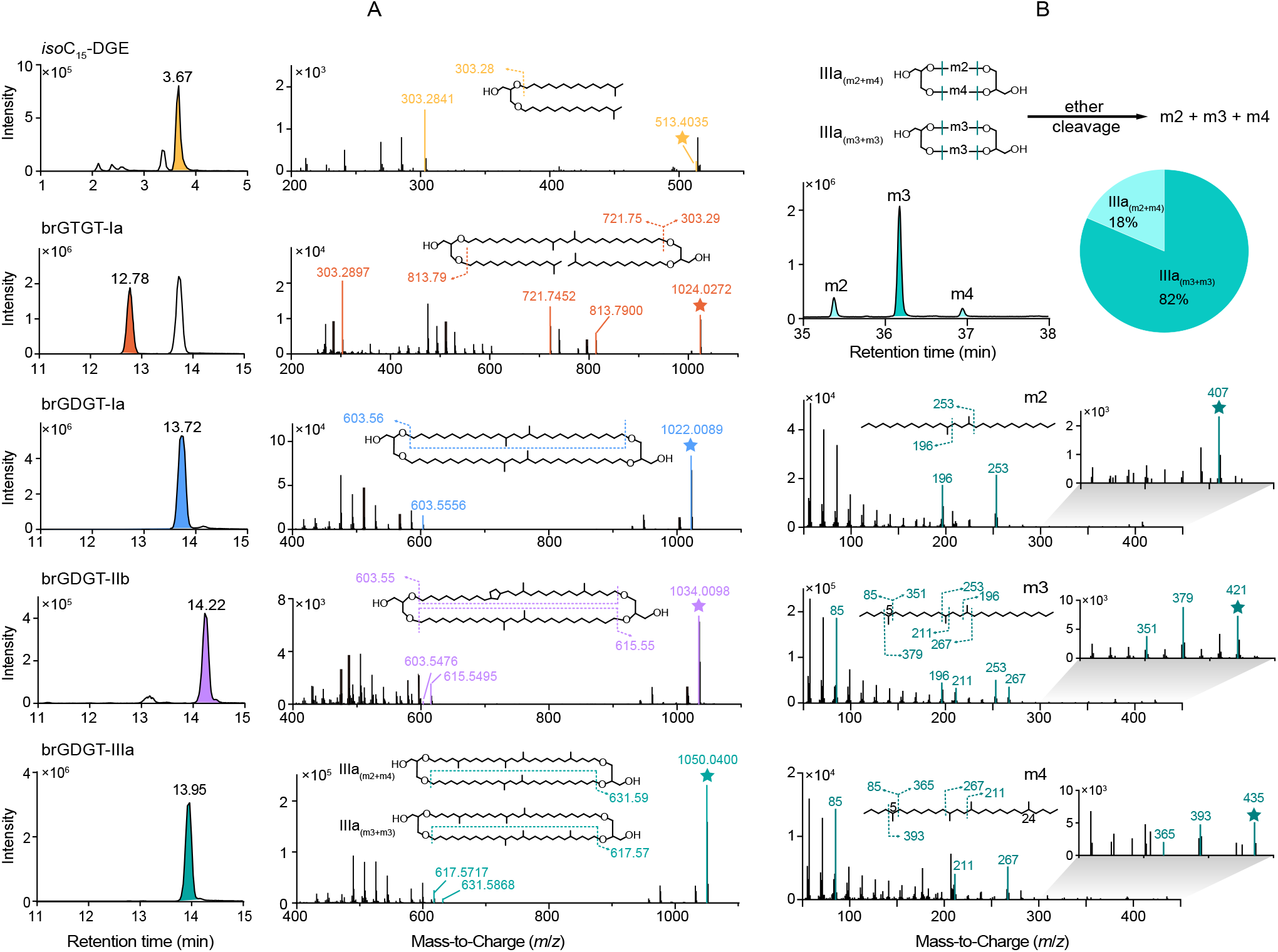
Identification of brGDGTs in strain Ellin6076 by RP−LC−HRMS and GC−MS. *(A)* Extracted ion chromatograms (EIC) and MS^2^ mass spectra of five representative compounds analyzed by RP−LC−HRMS are shown. *(B)* The partial total ion chromatogram (TIC) and mass spectra of brGDGT-IIIa derived alkyl chains analyzed by GC−MS are shown. IIIa_(m2+m4)_ and IIIa_(m3+m3)_ represent co-eluting isomers of brGDGT-IIIa with distinct alkyl chains. The m2, m3, and m4 refer to the alkyl chains with 2, 3, and 4 methyl groups, respectively. The precursor ions, such as [M+H]^+^ in RP−LC−HRMS and [M^+^−15] in GC−MS, are marked with stars. The characteristic product ions are marked by colors and fragment positions are denoted by dash lines. The pie chart shows the proportion of IIIa isomers in total IIIa. The calculation is based on the results of GC analysis and the contribution of co-eluting IIa to m2 and m3 in Fig. 1B is subtracted as described in *SI Appendix*.

The harvested cell mass was treated with acid hydrolysis to increase the yield of core lipids for the analyses of brGDGTs and their core lipid derivatives. The fragmentation patterns of five representative compounds are shown as examples in Fig. 1A. The MS^2^ behavior of *iso*C_15_-DGE corresponds to the loss of one *iso*C_15_-alkyl chain, resulting in a product ion of *m*/*z* 303.28 ([C_18_H_38_O_3_+H]^+^). This characteristic fragment ion is also present in the MS^2^ spectra of brGTGT-Ia as reported by Halamka et al. (2021) (13). The MS^2^ spectrum of brGDGT-Ia exhibits a featured product ion of *m*/*z* 603.56 ([C_36_H_74_O_6_+H]^+^) and brGDGT-IIb exhibits an additional product ion of *m*/*z* 615.55 ([C_37_H_74_O_6_+H]^+^), both of which are regular fragments observed in the MS^2^ spectra of brGDGT compounds (23).

To further determine the alkyl chain structures of brGDGT-IIIa isomers, brGDGT-IIIa purified from the total lipid extract of the hydrolyzed cells cultured at 10 °C was subjected to ether cleavage. The released alkanes were analyzed by gas chromatography–mass spectrometer (GC–MS). Three alkanes, including 13,16-dimethyloctacosane (m2), 5,13,16-trimethyloctacosane (m3), and 5,13,16,24-tetramethyloctacosane (m4), were found, indicating that brGDGT-IIIa was composed of two co-eluting isomers, i.e., one consisting of m2 and m4, and the other consisting of two m3 (Fig. 1B). The alkyl chain structures of the two brGDGT-IIIa isomers can also be confirmed by the fragment ions of *m*/*z* 631.59 ([C_38_H_78_O_6_+H]^+^) and *m*/*z* 617.57 ([C_37_H_76_O_6_+H]^+^) in the mass spectra of IIIa, which correspond to a neutral loss of m2 and m3 in IIIa_(m2+m4)_ and IIIa_(m3+m3)_, respectively (Fig. 1A). The estimated ratio of 82%:18% between %IIIa_(m3+m3)_ and %IIIa_(m2+m4)_ (the abundance percentage of IIIa isomers in total IIIa) suggests more IIIa_(m3+m3)_ was produced in the culture of strain Ellin6076, consistent with the observation in a peat sample (24).

The brGDGTs found in the environment contain isomers with an outer methyl group at either the α/ω5 or α/ω6 position, i.e. C_5_-methylated and C_6_-methylated brGDGTs (2). To determine which brGDGT isomers are produced by strain Ellin6076, we used a soil sample containing both C_5_- and C_6_-methylated brGDGTs as a reference, and compared the chromatogram and the retention time of target compounds using NP–LC–MS. The results showed that strain Ellin6076 produced C_5_-methylated brGDGTs (*SI Appendix*, Fig. S4). This is also confirmed by the GC−MS analysis of alkyl chains released from the ether cleavage of purified brGDGT-IIIa (Fig. 1B) and brGDGT-IIa (*SI Appendix*, Fig. S5).

To analyze the head groups of brGDGTs, we extracted IPLs from harvested cell mass with a modified Bligh-Dyer method. Phosphohexose (PH) was the most common polar head group detected in the culture, and this has also been identified in peat samples (25). IPLs such as PH-*iso*C_15_-DGE, PH-brGDGT-Ia, and PH-brGDGT-Ia-PH were identified and confirmed by MS^2^ spectra (*SI Appendix* Fig. S3).

### The abundance of brGDGTs in Acidobacteria cells

To estimate the proportion of brGDGTs in the total lipids of strain Ellin6076, the core lipid inventory of Ellin6076 cells was analyzed by gas chromatography–mass spectrometer (GC–MS) and NP–LC– MS. Strain Ellin6076 contains a variety of lipids including fatty acids, brGDGTs, hopanoids, and 3-hydroxy fatty acids. The total abundance of regular fatty acids, including saturated and unsaturated C_15−20_ fatty acids, was 13.8 fg/cell, accounting for 30% of the total quantified lipids. On the other hand, the abundance of all brGDGTs, including brGTGTs, was 30.3 fg/cell, accounting for 66% of the total lipids (Fig. 2). Compared to fatty acids and brGDGTs, *iso*C_15_ glycerol ethers including *iso*C_15_-monoalkyl glycerol ether (MGE) and *iso*C_15_-DGE had a much lower abundance, 0.9–1.8 fg/cell. Other lipids such as 3-hydroxy fatty acids and hopanoids were minor, with the summed abundance < 0.5 fg/cell (*SI Appendix*, Table S1). Intriguingly, *iso*-diabolic acids were absent from the lipid profile of strain Ellin6076. The fractional abundance of brGDGTs in the total lipids of strain Ellin6076 is much higher than that in *E. aggregans*, whose brGDGTs account for only approximately 3% of total lipids (13). Our findings demonstrate that some Acidobacteria, such as Ellin6076, use the membrane-spanning lipids brGDGTs and fatty acids as major components to form unique cell membranes with a mixed monolayer and bilayer structure (Fig. 2C).

**Fig. 2.**
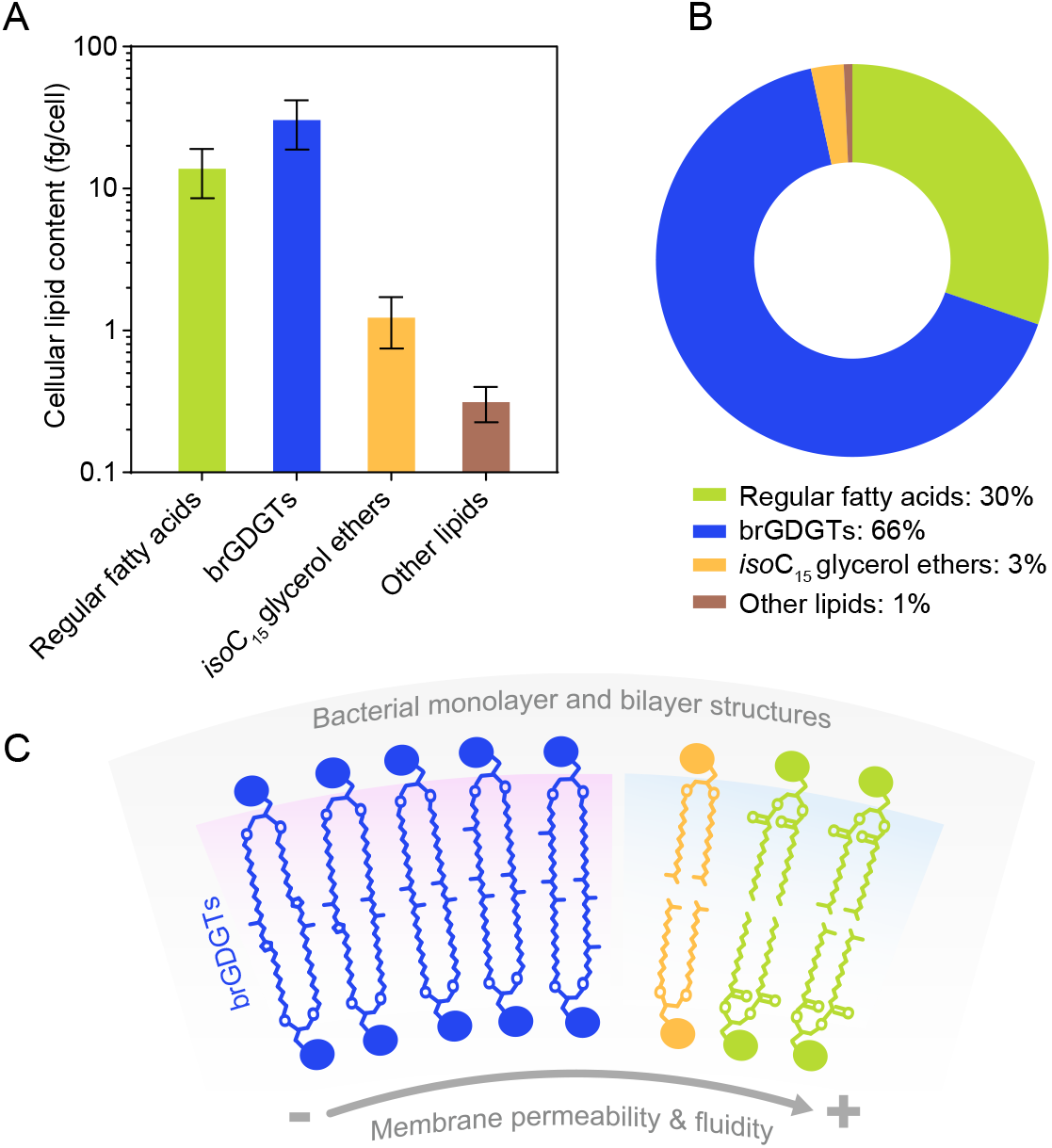
The lipid profile of strain Ellin6076. *(A)* The cellular contents and *(B)* the fractional abundances of major lipids in strain Ellin6076. The cellular content of brGDGTs includes that of brGTGTs, and the cellular content of *iso*C_15_ glycerol ethers includes that of *iso*C_15_-MGE and *iso*C_15_-DGE (*SI Appendix*, Table S1). Error bars represent the standard deviations among mean values of biological triplicate. The quantification is based on internal standards and cell amounts as described in *SI Appendix*. (*C*) Schematic cell membrane of strain Ellin6076 containing monolayer and bilayer structures with proposed permeability trend.

### The response of brGDGTs to cultivation conditions in Acidobacteria culture

The production of multiple brGDGTs by the strain Ellin6076 allows us to directly assess the brGDGT changes under controlled experimental conditions. We cultured strain Ellin6076 independently at temperatures ranging from 10–35 °C and pH ranging from 4.5–6.5 (*SI Appendix*, Fig. S6). Then we evaluated the changes in the fractional abundance of brGDGT-Ia, Ib, IIa, IIb, and IIIa, since these components are critical for the calculation of brGDGT-based proxies (e.g. MBT’_5ME_ and CBT_5ME_).

The MBT’_5ME_ index, expressing the methylation degree of C_5_-methylated brGDGTs showed a significant positive correlation with culture temperature (10 °C to 25 °C), having a determination coefficient (*R*^2^) up to 0.97 (Fig. 3A). Specifically, %Ia, the abundance percentage of brGDGT-Ia in total brGDGTs, increased with temperature in this range, while %IIa and %IIIa decreased (*SI Appendix*, Fig. S7, and Table S2). Importantly, the MBT’_5ME_ values at 30 °C and 35 °C were nearly 1.00 (*SI Appendix*, Table S2), the upper limit of this index. BrGDGT-Ia was overwhelmingly dominant at 30 °C (%Ia > 93%) and 35 °C (%Ia > 97%), consistent with environmental studies and confirming that the MBT’_5ME_ index is insensitive to temperature changes above 25 °C. Excluding these data showed a remarkable influence of temperature on the distributions of C_5_-methylated brGDGTs and the MBT’_5ME_ index in the culture of strain Ellin6076.

**Fig. 3.**
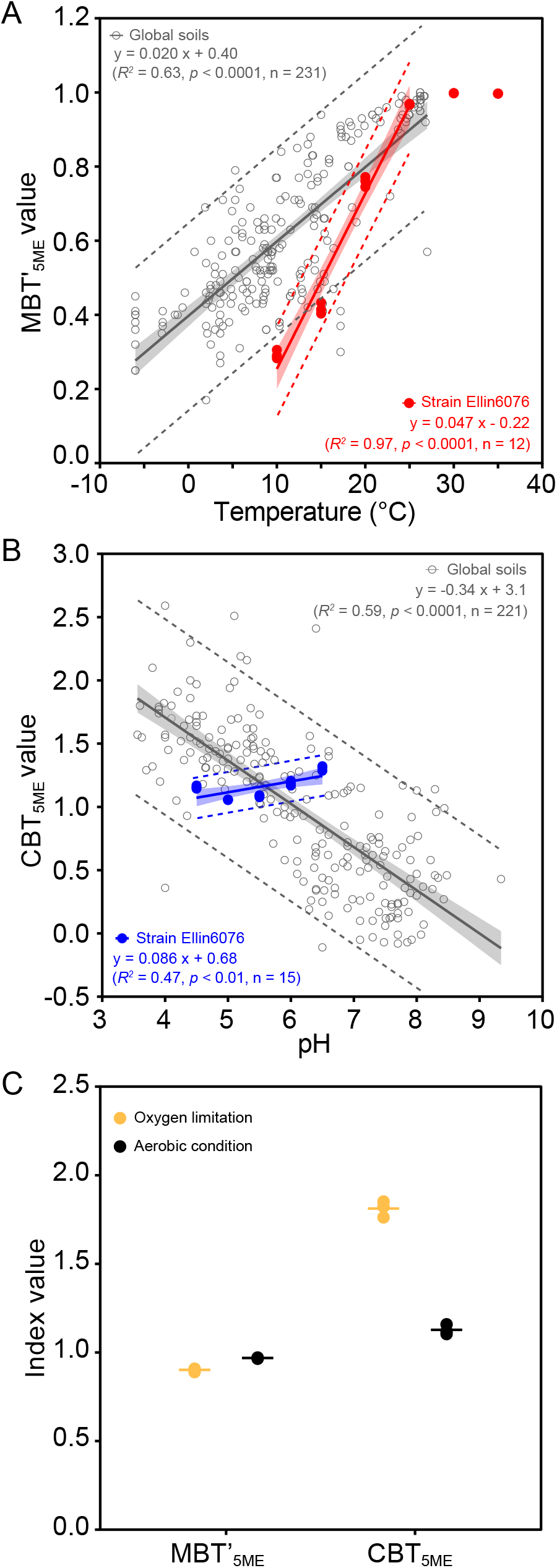
Relationships between brGDGT-based proxies and environmental factors. *(A)* The MBT’_5ME_ vs. temperature in the culture of strain Ellin6076 (red line) together with the MBT’_5ME_ in the global soil database (gray line; De Jonge et al., 2014). The data at 30 °C and 35 °C are excluded from the linear regression. *(B)* The CBT_5ME_ vs. pH in the culture of strain Ellin6076 (blue line) and the global soil database (gray line; De Jonge et al., 2014). The shaded area and dash lines show 95% confidence interval and 95% prediction interval of the linear regression, respectively. *(C)* The comparison of MBT’_5ME_ and CBT_5ME_ values between oxygen limitation and aerobic condition. Each biological replicate is displayed and lines represent the mean values.

The cyclization degree of brGDGTs was assessed by the CBT index, with a higher CBT value indicating a lower degree of cyclization (7). A positive correlation between CBT_5ME_ and pH from pH 4.5 to 6.5 was found for the culture samples, opposed to the general negative relationship derived from the global soil database (2, 7) (Fig. 3B).

In addition to temperature and pH, we also examined whether strain Ellin6076 adjusts its brGDGT composition in response to oxygen limitation. When strain Ellin6076 was cultured under oxygen limitation (1% O_2_ concentration, 25 °C, pH 5.5), the abundance of total brGDGTs decreased to 10 fg/cell compared to 30 fg/cell under aerobic condition (21% O_2_ concentration, 25 °C, pH 5.5), in contrast to the enhanced production under low oxygen reported in *E. aggregans* (13). Intriguingly, %IIa and %IIIa increased and %Ib, %Ic, and %IIb decreased under the stress of low oxygen. %Ia was effectively constant (87–91%) under both conditions (*SI Appendix*, Fig. S9 and Table S2). Consequently, MBT’_5ME_ values slightly decreased but CBT_5ME_ values strikingly increased under oxygen limitation (Fig. 3C).

## Discussion

### The conundrum of brGDGT-producers

The biological sources of brGDGTs in the environment have puzzled many scientists for more than two decades. Despite substantial effort devoted to solving this conundrum (12, 13, 16, 18–20, 26–28), a bacterial pure culture producing the multiple brGDGTs that have been used to construct paleoclimate has been lacking. Here, we found a bacterial strain Ellin6076 is capable of synthesizing multiple brGDGTs as its major cell membrane lipids. This is a crucial step towards understanding the source of diverse brGDGTs in the environment via culturing under different conditions, shedding new insights into the ecophysiology and taxonomy of brGDGT-producers.

Several lines of evidence, in particular from peats, suggest that brGDGT-producers might be anaerobic or facultative anaerobic bacteria (15, 18, 27). However, strain Ellin6076 is an obligately aerobic bacteria, indicating that at least a portion of brGDGTs in peats, soils, and lakes can be produced by aerobic Acidobacteria. This is consistent with one order of magnitude higher rates of brGDGT production in incubations with surface peat under oxic conditions than anoxic deeper peats (27), as well as the occurrence of abundant *in situ* brGDGTs produced in oxic lake water columns (29). However, these observations are inconsistent with the increased production of brGDGTs by *E. aggregans* under low O_2_ concentrations (13) and the high abundance of brGDGTs in low oxygen environments (15, 18). It is likely that brGDGTs are synthesized by a range of bacteria and/or at least some observations from the environment reflect preservation of brGDGTs in anoxic settings rather than higher production. Strain Ellin6076 uses glucose as the carbon source for the chemoheterotrophic lifestyle, consistent with the previous views on the lifestyle of brGDGT-producers based on the carbon isotopic compositions of brGDGTs in the environment (27, 30).

The previous identification of brGDGT-Ia in two Acidobacteria belonging to subdivision 1, together with the occurrence of more diverse brGDGTs in Ellin6076, indicates different Acidobacteria can produce completely distinct brGDGT profiles. This suggests that at least some variations in environmental brGDGT distributions reflect community change rather than physiological adaptations within a single taxon. In altitudinal or latitudinal transects with a large temperature or pH gradient, the impact of these environmental factors on brGDGTs overwhelmingly exceeds the community effect, resulting in significant correlations between brGDGT distribution and temperature or pH (9, 31, 32); however, at local scales, the community effect can dominate, as De Jonge et al. previously observed in high and mid-latitude soils (33, 34). It is, therefore, necessary to evaluate the community effect on existing brGDGT paleoclimate proxies in their applications to paleo-reconstructions.

For example, the occurrence of tetramethylated brGDGTs and C_5_-methylated brGDGTs with an absence of C_6_-methylated brGDGTs in Ellin6076 suggests that C_5_- and C_6_-methylated brGDGTs are produced by different (Acido)bacteria. The relative abundance of C_6_- vs. C_5_-methylated brGDGTs (generally expressed in the IR_6ME_ proxy) appears to be dependent on pH or salinity in a variety of environmental samples (10, 35, 36). This study suggests that pH proxies based upon the relative abundance of C_6_- vs. C_5_-methylated brGDGTs are essentially regulated by a shift in the Acidobacteria community. Ellin6076 falls within Acidobacteria subdivision 3, a clade that is abundant in acidic soils and peats (37–39), agreeing well with the dominance of C_5_-methylated brGDGTs over their C_6_-methylated isomers in these environments (9, 10). Future work should ascertain the biological source(s) of C_6_-methylated brGDGTs, perhaps by examining Acidobacteria with a Tes homolog that are abundant in alkaline environments, e.g. Acidobacteria subdivision 4 and 6.

### The physiological function of methylation and cyclization in brGDGTs

Our strain Ellin6076 culture experiments allow direct examination of how methylation and cyclization of brGDGTs in a single species respond to different temperature, pH and oxygen conditions. Canonically, modifications in the degree of methylation and cyclization are thought to be a microbial strategy – homeoviscous or homeostatic adaptation – to adapt to ambient environmental change (40, 41). Microbes modulate their lipid compositions to maintain appropriate fluidity and permeability of cell membranes (42, 43). Bacteria can modify the degree of branching in their fatty acids at varying temperatures (44), for example, with more branched-chain fatty acids observed at 45 °C than at 65 °C in the culture of a thermophilic bacteria *Bacillus stearothermophilus* (45). We observed a similar modification in Ellin6076, which produced more C_5_-methylated brGDGTs (e.g. brGDGT-IIa and IIIa) and fewer tetramethylated brGDGT-Ia, thereby increasing the degree of methylation, at temperatures below 25 °C. Recent molecular dynamics simulations of bacterial membranes consisting of brGDGTs confirm that a higher degree of methylation results in a less rigid and more fluid membrane (46). Our culturing experiments support this theory and suggest that the increase in brGDGT methylation is a physiological adaptation strategy for brGDGT-producing bacteria in cold conditions. In contrast, pH and oxygen limitation exert a minor effect on the degree of brGDGT methylation (*SI Appendix*, Fig. S8 and Fig. S9).

The degree of cyclization in isoGDGTs is a key strategy for archaea to adapt to extreme environments, with more cyclopentyl moieties generally produced by archaea growing at a higher temperature or a lower pH (47–49). Molecular modeling suggests that an increase in isoGDGT cyclization degree leads to tighter membrane packing that enhances membrane thermal stability and reduces overall membrane permeability (50, 51). Given the similarity in structure, it is reasonable to hypothesize that cyclopentane rings in brGDGTs have a similar function. Indeed, our results demonstrate that lower pH and higher temperature generally cause a higher degree of brGDGT cyclization (*SI Appendix*, Fig. S7 and Fig. S8), which is consistent with the behaviour of isoGDGTs in a thermoacidophilic archaeon *Sulfolobus acidocaldarius* (49). However, this relationship is opposite to the well-established empirical relationship of soil pH with the degree of brGDGT cyclization. This could mean that Ellin6076 (or our culture conditions) are atypical or that brGDGT cyclization is sensitive to multiple variables, instead of pH alone. Given the similarity in behaviour between the cyclization of Ellin6076 brGDGTs and that in archaea, as well as the predictions of molecular modelling, we instead propose that the widely observed environmental relationship documents changes in the brGDGT-producing community rather than an ecophysiological relationship.

### Implications for brGDGT-based temperature and pH proxies

The physiological function of methylation and cyclization in brGDGTs helps to interpret MBT and CBT proxies and their relationships with environmental factors in nature. The positive correlation between MBT’_5ME_ and temperature for the culture of Ellin6076 is consistent with the empirical observation of global soils (Fig. 3A). The calibration equation for the strain Ellin6076 is:

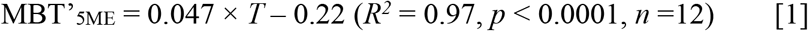

The slope of the regression line for strain Ellin6076 is significantly steeper than that for the global soil dataset. There could be two explanations for this. First, the majority of brGDGTs in soils are produced by other (Acido)bacteria species with a brGDGT response to temperatures differing from the strain Ellin6076. Second, the MBT’_5ME_ for global soils could record the growing season temperatures. The difference in MBT’_5ME_ values between the strain Ellin6076 and global soils increases with decreasing temperature, which might relate to the increased seasonal production of brGDGTs in colder soils. The strain Ellin6076 cannot grow well below 10 °C. Likewise, Acidobacteria producing brGDGTs in soils are unlikely to proliferate at a low temperature. If the MBT’_5ME_-temperature calibration for the strain Ellin6076 is applied to the global soils, the temperature estimates for soils from cold regions would be significantly higher than the mean annual air temperature (MAT). In contrast, the MBT’_5ME_ values for the strain Ellin6076 agree well with those for global soils at temperatures > 20 °C, where the effect of temperature seasonality is minor. This suggests caution in the application of brGDGT temperature proxies in low temperature contexts. At the same time, MBT’_5ME_ values reached saturation at 25 °C in the culture experiments (*SI Appendix*, Table S2), which is consistent with the observations in soils, suggesting the MBT’_5ME_ index cannot be used to reconstruct temperature changes above that (Fig. 3A).

Temperature was the only factor controlling MBT’_5ME_ variation in the culture experiment, whereas other environmental variables such as pH and oxygen availability exerted only minor effects on this index. The MBT’_5ME_ values for strain Ellin6076 barely changed under different pH conditions (*SI Appendix*, Table S2) and decreased only slightly under oxygen limitation (Fig. 3C). This suggests that pH and oxygen availability could be excluded as potential factors affecting the MBT’_5ME_ in the environment. However, this only applies at the species level. Several previous studies showed that soil pH or oxygen level are key factors that determine the variation of MBT’_5ME_ in soils and peats (10, 52, 53). We attribute this to the community effect inferred above. A shift in the Acidobacteria community would cause a change in brGDGT distribution since some Acidobacteria can only produce brGDGT-Ia (12, 13), whereas other Acidobacteria like Ellin6076 are capable of synthesizing more diverse brGDGTs. It also appears that changes in pH drive shifts in the Acidobacteria community as documented by the unexpected change in brGDGT cyclization (see above). This could explain why changes in brGDGT-reconstructed pH are sometimes associated with unexpected changes in MBT-derived temperatures (54, 55).

In addition, oxygen limitation could trigger the production of more brGDGTs, especially brGDGT-Ia (12, 13). Although not observed in strain Ellin6076, this effect would also increase the MBT’_5ME_ values in soils. While our study demonstrates the robustness of MBT’_5ME_ as a paleothermometer, factors that potentially affect the Acidobacteria community (not limited to oxygen limitation and pH) need to be considered as well.

Our work provides fundamental new insights into brGDGT biosynthesis and adaptation. By identifying archaeal tetraether synthase homologs in bacterial genomes, we were able to identify a brGDGT-producing Acidobacterium, strain Ellin6076. Crucially, because this organism shows to produce a suite of brGDGTs, we were able to conduct culture experiments to determine the response of brGDGT distributions to changes in temperature, pH and oxygen. Such work reaffirms confidence in brGDGT-temperature proxies but also suggests that much of the environmental variation in brGDGT distributions ascribed to pH is instead due to community change.

## Methods

### Strain and Culturing

*Candidatus Solibacter usitatus* Ellin6076 (DSM 22595) was purchased from the German Collection of Microorganisms and Cell Cultures (DSMZ). The strain was routinely cultured in the modified liquid MM medium under the optimal growth conditions aerobically at 25 °C and pH 5.5 for 14 days to reach the stationary phase. To investigate how brGDGT distribution responds to environmental changes, strain Ellin6076 was cultured under different temperatures (10−35 °C), pH values (4.5−6.5), and oxygen levels (1% vs. 21% O_2_ concentration in the headspace). All the experiments were performed in biological triplicates. The details of culturing experiments are described in *SI Appendix*.

### Lipid Extraction and Analyses

Culture samples were harvested at the stationary phase and collected by 10,000 × g centrifugation for 15 min and cell pellets were kept at –80 °C before further experiments. An aliquot of wet cell mass was treated with acid hydrolysis for core lipid (CL) analysis, and another aliquot was directly extracted using a modified Bligh-Dyer method for intact polar lipid (IPL) analysis as described in *SI Appendix*.

BrGDGTs together with their CL and IPL derivatives were identified using a Waters ACQUITY I-Class Ultra-performance liquid chromatography (UPLC) coupled to SYNAPT G2-S*i* quadrupole time-of-flight (qTOF) high-resolution mass spectrometer. The quantification of brGDGTs with the C_46_ GTGT internal standard was performed on an Agilent 1260 series high-performance liquid chromatography (HPLC) system coupled with Agilent 6135B quadrupole mass spectrometer. The CL inventory including fatty acids, mono/dialkyl glycerol ethers, and other lipids in the cultures was analyzed by a Thermo Finnigan Trace 1300 gas chromatography coupled to an ISQ 7000 mass spectrometer (GC−MS). The purification and ether cleavage were performed on target compounds such as brGDGT-IIa and IIIa to further determine the methyl positions of brGDGTs and co-eluting isomers of IIIa produced by strain Ellin6076. The alkanes released from brGDGTs were analyzed by GC and GC−MS. The details of lipid extraction, analyses, identification and quantification are described in *SI Appendix*.

### Calculation of MBT and CBT proxies

We used MBT’_5ME_ and CBT_5ME_ proxies following De Jonge et al. (2014) to evaluate the distribution of brGDGTs (2), as only C_5_-methyl isomers were identified in strain Ellin6076. The calculation was based on the relative abundances of major brGDGTs:

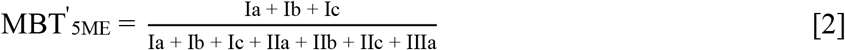

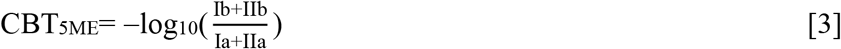

## Acknowledgments

We thank Wenyong Yao, Wan Zhang, and Jing Guo for microscopy observation and cell counting. This study was supported by the National Natural Science Foundation of China (No. 92051112 to Z.Z., No. 32170041 to Z.Z., and No. 42073072 to H.Y.), the Science, Technology, and Innovation Commission of Shenzhen Municipality (No. 20200925154325002 to Z.Z.), the Southern Marine Science and Engineering Guangdong Laboratory (Guangzhou) (No. K19313901), and the Shenzhen Key Laboratory of Marine Archaea Geo-Omics, Southern University of Science and Technology (No. ZDSYS201802081843490).

## Notes

### Competing Interest Statement

The authors have declared no competing interest.

