## Supplementary Information for "The production of diverse brGDGTs by an Acidobacterium allows a direct test of temperature and pH controls on their distribution"

**This PDF file includes:**

Supplementary text
Figs. S1 to S9
Tables S1 to S2
SI Reference

### Supplementary Information Text

### Methods

**Strain and culturing.** Acidobacterial strain *Candidatus Solibacter usitatus* Ellin6076 (DSM 22595) was purchased from the German Collection of Microorganisms and Cell Cultures (DSMZ). The strain was cultured in liquid MM medium with slight modifications, containing $KH_2PO_4$ 0.1 g/l, $(NH_4)_2SO_4$ 0.2 g/l, $MgSO_4 \cdot 7H_2O$ 0.1 g/l, $CaCl_2 \cdot 2H_2O$ 0.02 g/l, yeast extract 0.2 g/l and glucose 3 mM at final concentration (1). Trace element solution SL10, selenite-tungstate solution, and vitamin solution were supplemented following the description in VL55 medium (https://www.dsmz.de/microorganisms/medium/pdf/DSMZ_Medium1266.pdf). Medium was buffered by 10 mM 2-morpholinoethanesulfonic acid (MES) at the final concentration and adjusted to pH 5.5 using NaOH/KOH solution. Cultures were kept shaking at 200 RPM in the dark (2). The strain was routinely cultured under the optimal growth conditions aerobically at 25 °C and pH 5.5 for 14 days to reach the stationary phase. All the experiments were performed in biological triplicates. Optical density at 600 nm ($OD_{600}$) was employed to routinely monitor the growth. The specific growth rate was calculated by the logarithmic transformation of measured $OD_{600}$ values and linear regression fitting (3).

We performed independently culturing experiments to evaluate brGDGTs response of strain Ellin6076 to the following parameters: temperature (10–35 °C), pH (4.5–6.5), and oxygen level (1% vs. 21% $O_2$ concentration in the headspace). For the temperature experiment, cultures were aerobically grown in triplicate at pH 5.5 and temperatures of 10, 15, 20, 25, 30, and 35 °C, respectively. For the pH experiment, cultures were aerobically grown in triplicate at 25 °C with pH of 4.5, 5.0, 5.5, 6.0, and 6.5, respectively. A 30 mM MES buffer at final concentration was used to maintain the designed pH conditions. The initial $OD_{600}$ values were approximately 0.02–0.03 at the beginning of per experiment. Culture samples were harvested at the stationary phase and collected by 10,000 × g centrifugation for 15 min. Culture samples at 35 °C were collected using 0.22 μm polyvinylidene fluoride (PVDF) filters (Millipore) due to the low cell biomass. The cell pellets and filters were subsequently kept at –80 °C before further experiments. OD measurements were not available to assess the growth below 20 °C as strain Ellin6076 grew very slowly and flocculated into visible aggregates. Therefore, culture samples at 10 °C and 15 °C were collected after 40 days and 26 days, respectively, when the culture became turbid and apparently aggregated.

The experiment of oxygen limitation was carried out as previously described (3). The 250 ml serum bottles containing 100 ml medium were sealed with air-tight butyl rubber septa and caps and continuously stirred and flushed with sterile nitrogen to remove oxygen of the 150 ml-headspace. Then the bottles including the medium therein were autoclaved and supplemented with filtered solutions of glucose, trace elements, selenite-tungstate and vitamins. Another 7.5 ml of sterile air was injected to achieve 1% $O_2$ concentration in the 150 ml-headspace before the inoculation, leading to the oxygen limitation compared to the aerobic condition (21% $O_2$ concentration). The cultures were grown in triplicate under oxygen limitation and aerobic condition, respectively. Other parameters were kept as the optimal conditions (25 °C, pH 5.5, shaking at 200 RPM in the dark). OD measurements were not available to assess the growth due to the aggregates under the oxygen limitation. Samples were collected after 14-day incubation using the above protocol.

**Lipid extraction.** The biomass for core lipid (CL) and intact polar lipid (IPL) analyses was both approximately $1\times10^8$–$1\times10^9$ cells. To obtain the hydrolyzed total lipid extracts

(H-TLEs) for CL analysis, an aliquot of wet cell pellets was subjected to acid hydrolysis following the protocol in Zeng et al. (2019) (4). Briefly, cells were resuspended with 1 ml ultrapure water, and a 50 μl $C_{46}$ GTGT internal standard (concentration of 1.189 ng/μl) was added. Then the cells were hydrolyzed with 5−10 ml methanol (MeOH): hydrochloric acid (HCl) = 90:10 (v:v) at 70 °C for 8 hrs. After hydrolysis, 10 ml ultrapure water and 10 ml dichloromethane (DCM) were added for the separation of aqueous and organic phases. The bottom layer of DCM was transferred and collected. The hydrolyzed liquid was additionally extracted twice with fresh DCM. All DCM fractions were combined and filtered through a 0.45-μm polytetrafluoroethylene (PTFE) filter to form H-TLEs for CL detection, which were then dried under a gentle stream of nitrogen gas and stored at –80 °C before further analysis.

Another aliquot of wet cell pellets was directly extracted using a modified Bligh-Dyer method described in Chen et al. (2018) for the IPL analysis (5). A 50 μl $C_{46}$ internal standard (concentration of 1.189 ng/μL) was added and cell pellets were extracted twice via sonication in 9.5 ml mixture of MeOH : DCM : phosphate buffer (PB, pH 7.4) = 2:1:0.8 (v:v:v) for 15 min and the supernatants were collected after centrifugation. The 19 ml combined supernatants were then adjusted to the solvent ratio of MeOH : DCM : PB = 1:1:0.9 (v:v:v) for the phase separation and the DCM layer was collected. The remaining liquid was additionally extracted twice with fresh DCM. All organic phases were combined to yield non-hydrolyzed TLEs containing both CLs and IPLs, which were then dried under a gentle nitrogen stream and stored at –80 °C before further analysis.

**BrGDGT analyses.** H-TLEs were analyzed with normal-phase–liquid chromatography–mass spectrometry (NP–LC–MS) to obtain the distribution of brGDGTs in the culture. An Agilent 1260 series high-performance liquid chromatography (HPLC) system coupled with Agilent 6135B quadrupole mass spectrometer was employed and operated at positive ionization mode with an atmospheric pressure chemical ionization (APCI) source. BrGDGTs were separated using two BEH HILIC columns (2.1 × 150 mm, 1.7 μm; Waters) in tandem as described in Hopmans et al. (2016) (6). The flow rate was 0.2 ml/min and columns were kept at 30 °C. The total run time was 120 min and the injection volume was 10 μl for each sample. The elution gradient was set following Hopmans et al. (2016) with solvent A of *n*-hexane and solvent B of *n*-hexane: *iso*-propanol = 9:1 (v:v). Compounds were eluted with 18% B in the first 25 min, and then linearly increased to 35% B in the next 25 min, and subsequently 100% B in another 30 min. Finally, the gradient returned to 18 B% in 10 min and re-equilibrated for 30 min. MS condition was identical to Schouten et al. (2007): Nebulizer pressure 60 psi, vaporizer temperature 400 °C, drying gas (nitrogen) flow 6 l/min at 200 °C, capillary voltage 3 kV, corona 5 μA (7). BrGDGTs and several CL derivatives were scanned using selective ion monitoring (SIM) mode to increase the sensitivity (*m*/*z*: 513.5, 743.6, 1018, 1020, 1022, 1024, 1032, 1034, 1036, 1038, 1046, 1048, 1050, 1052). The diagnostic soil samples were analyzed in the same batch as references for compound identification. Lipids were identified with the retention time and mass spectra of protonated ions ($[M+H]^+$).

Non-hydrolyzed TLEs as well H-TLEs of the cultures grown at 10 °C and 25 °C were analyzed with reverse-phase–liquid chromatography–high-resolution mass spectrometry (RP–LC–HRMS) for structure identification. A Waters ACQUITY I-Class Ultra-performance liquid chromatography (UPLC) coupled to SYNAPT G2-S*i* quadrupole time-of-flight (qTOF) high-resolution mass spectrometer with electrospray ionization (ESI) source in positive mode was employed. Compounds were separated with a $C_{18}$-AR UPLC column (2.1 × 150 mm, 2 μm; ACE) maintained at 55 °C. The total run time was 40 min and samples were kept at 7 °C during the run. The flow rate was 0.3 ml/min. A total of 5 μl sample was injected and eluted with the following gradient: 100% A maintained for the first 5 min, the gradient of solvent B linear increased to 24% at 10 min, 49% at 28 min, 60% at 30 min, 90% at 32 min, followed by re-equilibration for 5 min. Solvent A was MeOH (Optima™ LC/MS Grade, Fisher Chemical) and Solvent B was *iso*-propanol (Optima™ LC/MS Grade, Fisher Chemical), both with 0.1% $NH_3OH$ (25–30% $NH_3$ basis, Sigma-Aldrich) and 0.04% formic acid ( > 99.0%, Optima™ LC/MS Grade, Fisher Chemical) (8). MS setting was as follows: Capillary 2.5 kV, source temperature 120 °C, sampling cone 45, source offset 80, desolvation gas flow 800 l/hr at 350 °C, cone gas flow 50 l/hr, nebulizer gas flow 6.5 bar.

The analyzer was performed at resolution mode. The mass acquisition mode was FAST-DDA with a mass range for MS *m/z* 100–2,000 and $MS^2$ *m/z* 50–2,000. The scan time for MS and $MS^2$ was both 0.2 sec. Top 5 ions of the highest intensity with a TIC threshold > 20,000 counts were fragmented via collision-induced dissolution (CID) to obtain product ion spectra. A real-time dynamic exclusion of masses (acquired and excluded for 5 sec) was enabled to acquire $MS^2$ spectra for fewer intensity ions. A ramped collision energy of the transfer cell was used to generate product ion spectra, which started at 10 V and ended at 55 V for low mass and start at 15 V and ended at 65 V for high mass, respectively. The accuracy of MS was calibrated at the beginning of the experiments using a solvent of sodium iodide (*m/z* 50–2,000; Residual mass error < 0.5 ppm). A leucine enkephalin solution ($[M+H]^+$ at *m/z* 556.2771) was employed as the lockmass reference compound to perform real-time calibration (scan time 0.2 sec, interval 20 sec).

**Fatty acid analysis.** Cell biomass of the *Ca. Solibacter usitatus* Ellin6076 cultured under the optimal growth conditions (25 °C, pH 5.5) was subjected to acid hydrolysis following the above protocol to obtain H-TLEs. The H-TLEs were spiked with a known amount of internal standard, cholanic acid, and esterified with $BF_3$/MeOH solution at 70 °C for 2 hrs. The fatty acid methyl esters (FAMEs) were recovered by extracting the solution with *n*-hexane (6×). Due to the occurrence of monoalkyl glycerol ethers (MGEs) and diethers, the resulting lipids were further derivatized with bis(trimethylsilyl)trifluoroacetamide (BSTFA) at 70 °C for 1.5 hrs and dried under a gentle stream of nitrogen gas. Lipids were analyzed by a Thermo Finnigan Trace 1300 gas chromatography coupled to an ISQ 7000 mass spectrometer (GC–MS) equipped with a DB-5MS silica capillary column (60 m × 0.25 mm × 0.25 μm) with helium as carrier gas. The oven temperature was programmed from 70 ℃ to 210 ℃ at 10 ℃/min and then to 310 ℃ at 3 ℃/min, with the final temperature being held at 310 ℃ for 26 min. The EI energy was 70 eV and the auxiliary temperature of GC and MS was 310 ℃.

**Purification and ether cleavage of brGDGT-IIa and IIIa.** To further determine the methyl positions of major brGDGTs and alkyl chain structures of brGDGT-IIIa isomers, brGDGT-IIa, and IIIa in the H-TLEs of cultures grown at 10 ℃ were purified by a semi-preparative HPLC on an Agilent series 1290 HPLC equipped with a fraction collector. Compound separation was achieved with an Alltech Prevail Cyano column (150 mm × 2.1 mm, 3 μm). The elution gradient followed Schouten et al. (2007) (7). GDGTs were first eluted isocratically for the first 5 min with 90% A and 10% B, followed by an increase to 18% B from 5 to 45 min, where A was *n*-hexane and B was *n*-hexane: *iso*-propanol = 9:1 (v:v). 100% B was then used to wash the column for the next 10 min.

The fractions containing brGDGT-IIa and IIIa were transferred to vials and dried under a gentle stream of nitrogen gas. Boron tribromide ($BBr_3$) (0.5 ml, 1 M in DCM; Sigma-Aldrich) was added to cleave the ether bonds of brGDGTs. The vial was sealed and heated at 60 °C for 2 hrs. After the residual solvents were removed in a gentle stream of nitrogen gas, the bromides were reduced to alkanes by superhydride (0.5 ml, lithium triethylborohydride ($LiEt_3BH$), in DCM, Sigma-Aldrich) in the sealed vial with an argon atmosphere at 60 °C for 2 hrs. To quench the reaction, a small amount of ultrapure water was added. The solution was extracted (6×) with *n*-hexane. The alkanes released from brGDGTs were analyzed by an Agilent 8890 GC and an Agilent 6890 GC coupled to a 5975 MS with the same GC and MS conditions as described above. Alkanes were identified based on the mass spectra as described in the previous study (9).

**Identification of brGDGTs.** Structural assignments of brGDGTs and their derivatives based on RP−LC−HRMS were achieved by the accurate molecular mass (mass error < 10 ppm) and their diagnostic product ions in $MS^2$ spectra. The derivatives identified in this study included *iso*$C_{15}$-dialkyl glycerol ether (DGE), branched glycerol trialkyl glycerol tetraethers (brGTGTs), branched glycerol dialkanol diethers (brGDDs) and IPLs corresponding to some above core lipids.

Protonated $[M+H]^+$, ammoniated $[M+NH_4]^+$ and sodiated $[M+Na]^+$ ions are common pseudo molecular ions observed for all compounds under RP−LC−HRMS. The characteristic product ions in $MS^2$ spectra of brGDGTs under CID are associated with neutral losses of one $H_2O$ molecule (i.e. $[M+H–18.01]^+$), one glycerol (i.e. $[M+H–74.04]^+$) and alkyl units as previously reported (10). The peak of *iso*$C_{15}$-DGE ($[M+H]^+$ at *m*/*z* 513.4944) was observed at 3.67 min. The $MS^2$ behavior of *iso*$C_{15}$-DGE corresponds to the loss of one *iso*$C_{15}$-alkyl chain ($C_{15}H_{30}$), resulting in a product ion of

$m/z$ 303.28. This fragment ion also occurs as a major product ion in $MS^2$ spectra of brGTGT-Ia ($[M+H]^+$ at $m/z$ 1024.0270) reported by Halamka et al. (2021) (3). The $MS^2$ spectra of brGDGT-Ia ($[M+H]^+$ at $m/z$ 1022.0093) exhibit a featured product ion of $m/z$ 603.56 by a loss of one alkyl chain and a series of subsequent loss of $H_2O$ and glycerol (10). The losses of alkyl chains of brGDGT-IIb ($[M+H]^+$ at $m/z$ 1034.0090) generate characteristic product ions of $m/z$ 615.55 and $m/z$ 603.55, revealing the additional methyl locating at the same side with the cyclopentyl ring (Fig. 1). The $MS^2$ spectra of other typical brGDGTs and brGTGTs were illustrated in *SI Appendix,* Fig. S1 and Fig. S2. BrGDGT-Ic and IIc were tentatively detected and assigned with the retention time by NP−LC−MS (*SI Appendix,* Fig. S4). The $MS^2$ spectra of these two compounds were not obtained, due to the low abundance of these compounds and relatively low sensitivity under full scan mode of RP−LC−HRMS. The structural identification of brGDDs followed the description in Liu et al., (2012) (10) and the fragmentation patterns were similar to those in brGDGTs (*SI Appendix,* Fig. S2).

Phosphohexose (PH) was the most common polar head group of brGDGTs detected in the non-hydrolyzed extracts. The PH-*iso*$C_{15}$-DGE was detected at 1.90 min and the $MS^2$ spectra were characterized by product ions of $m/z$ 593.49 (loss of one hexose) and $m/z$ 383.25 (loss of one hexose plus one *iso*$C_{15}$-alkyl chain) (*SI Appendix,* Fig. S3). BrGDGTs with one and two PH headgroups were detected at 8.81−8.94 min and 4.06–4.14 min, respectively. The product ion of $m/z$ 261.04 (PH headgroup) was commonly observed in these compounds that had similar fragmentation patterns usually associated with neutral losses of $H_2O$ and one hexose (11).

**Quantification and calculation.** The cell density of strain Ellin6076 was estimated based on the microscopic counts and measured $OD_{600}$ values. The cell density was approximately 1.68 × $10^8$ cell/ml at $OD_{600}$ = 0.46, when the strain was cultured to the stationary phase under the optimal growth condition. The cell amount for lipid extraction was then determined by the cell density at a specific $OD_{600}$ value and the volume of culture.

The abundances of brGDGTs were calculated by comparison of each brGDGT peak area with the peak area of internal standard $C_{46}$ GTGT based on the analysis of NP−LC−MS. The fractional abundance (%) of individual brGDGT compound was calculated via dividing the peak area of each brGDGT by the sum of the peak areas of all brGDGTs. The abundances of regular fatty acids, *iso*$C_{15}$ glycerol ethers (i.e. *iso*$C_{15}$-MGE and *iso*$C_{15}$-DGE), hopanoids, and 3-hydroxy fatty acids, were calculated by the comparison of the peak areas between target compounds and the internal standard, cholanic acid. The final abundances were normalized to cell amounts to obtain the cellular lipid contents (fg/cell) (12). The fractional abundances (%) of different lipid components, such as regular fatty acids, brGDGTs and *iso*$C_{15}$ glycerol ethers, were the percentages of the cellular content of each lipid component to the total lipids.

The proportion of brGDGT-$IIIa_{(m3+m3)}$ in total IIIa isomers was estimated based on the peak area of IIIa-released alkanes analyzed by GC. Since m2 and m4 were both released from $IIIa_{(m2+m4)}$, they should have equivalent abundances. However, we found that the peak area of m2 was almost 2 fold more than that of m4, indicating a small amount of IIa exists in the purified IIIa and causes higher abundances of m2 and m3 than expected. We subtracted the contribution of co-eluting IIa (the difference in peak area between m2 and m4) from the peak area of m2 and m3, respectively. We assumed the peak area of m4 to represent the abundance of $IIIa_{(m2+m4)}$, and half of the peak area of m3 after calibration to represent the abundance of $IIIa_{(m3+m3)}$. The abundance percentage (%) of each IIIa isomer in total IIIa isomers was calculated and the estimated ratio between %$IIIa_{(m3+m3)}$ and %$IIIa_{(m2+m4)}$ was 82%:18% in the culture of strain Ellin6076 at 10 °C.

We used $MBT'_{5ME}$ and $CBT_{5ME}$ indices following De Jonge et al. (2014) to evaluate the distribution of brGDGTs (13), as only $C_5$-methyl isomers were identified in the culture of strain Ellin6076. The calculation was as follows and the roman numerals below referred to the relative abundance of brGDGTs:

$$MBT'_{5ME} = \frac{Ia + Ib + Ic}{Ia + Ib + Ic + IIa + IIb + IIc + IIIa}$$

$$CBT_{5ME} = -\log_{10}\left(\frac{Ib+IIb}{Ia+IIa}\right)$$

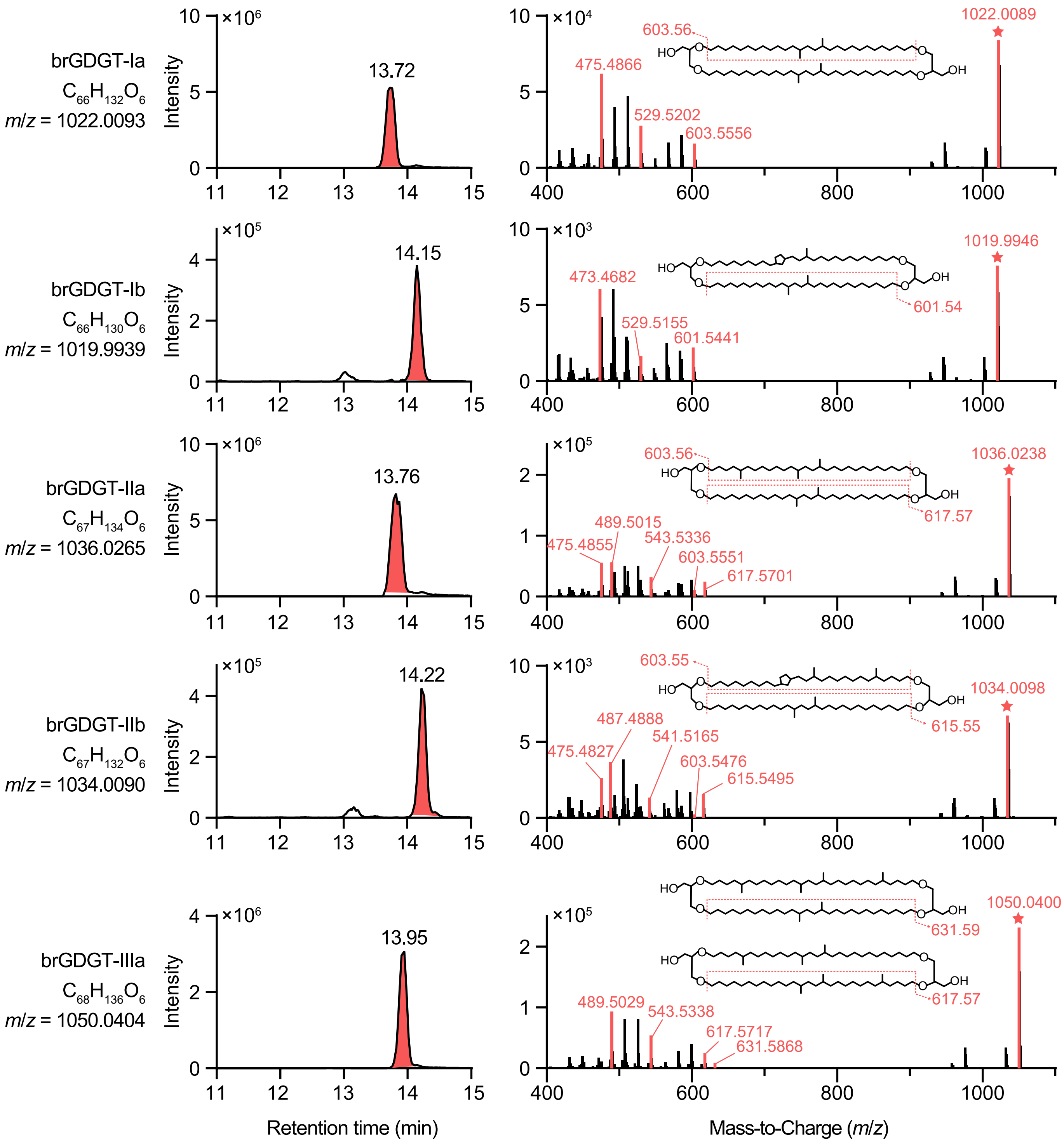


**Fig. S1**. Extracted ion chromatograms and MS² mass spectra of regular brGDGTs in the strain Ellin6076 analyzed by RP–LC–HRMS. The precursor ions ($[M+H]^+$) and characteristic product ions are marked by stars and red color, respectively, and fragment positions are denoted by dash lines. BrGDGT-Ic and IIc were tentatively detected and assigned with retention time by NP–LC–MS. The MS² spectra of these two compounds were not obtained, due to the low abundance of these compounds.

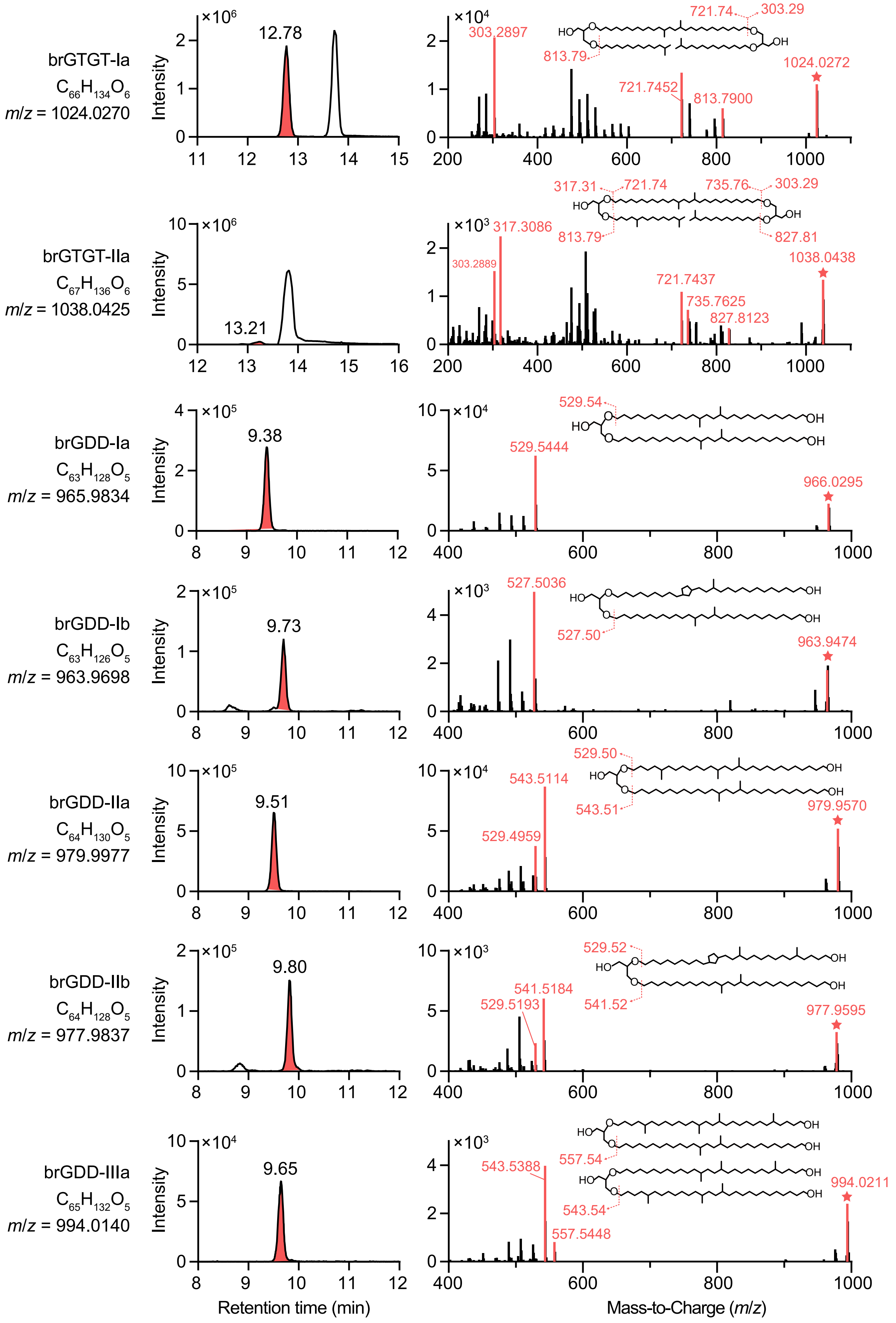


**Fig. S2.** Extracted ion chromatograms and MS[2] mass spectra of major brGTGTs and brGDDs identified in strain Ellin6076 using RP–LC–HRMS. The precursor ions ($[M+H]^+$) and characteristic product ions are marked by stars and red color, respectively, and fragment positions are denoted by dash lines.

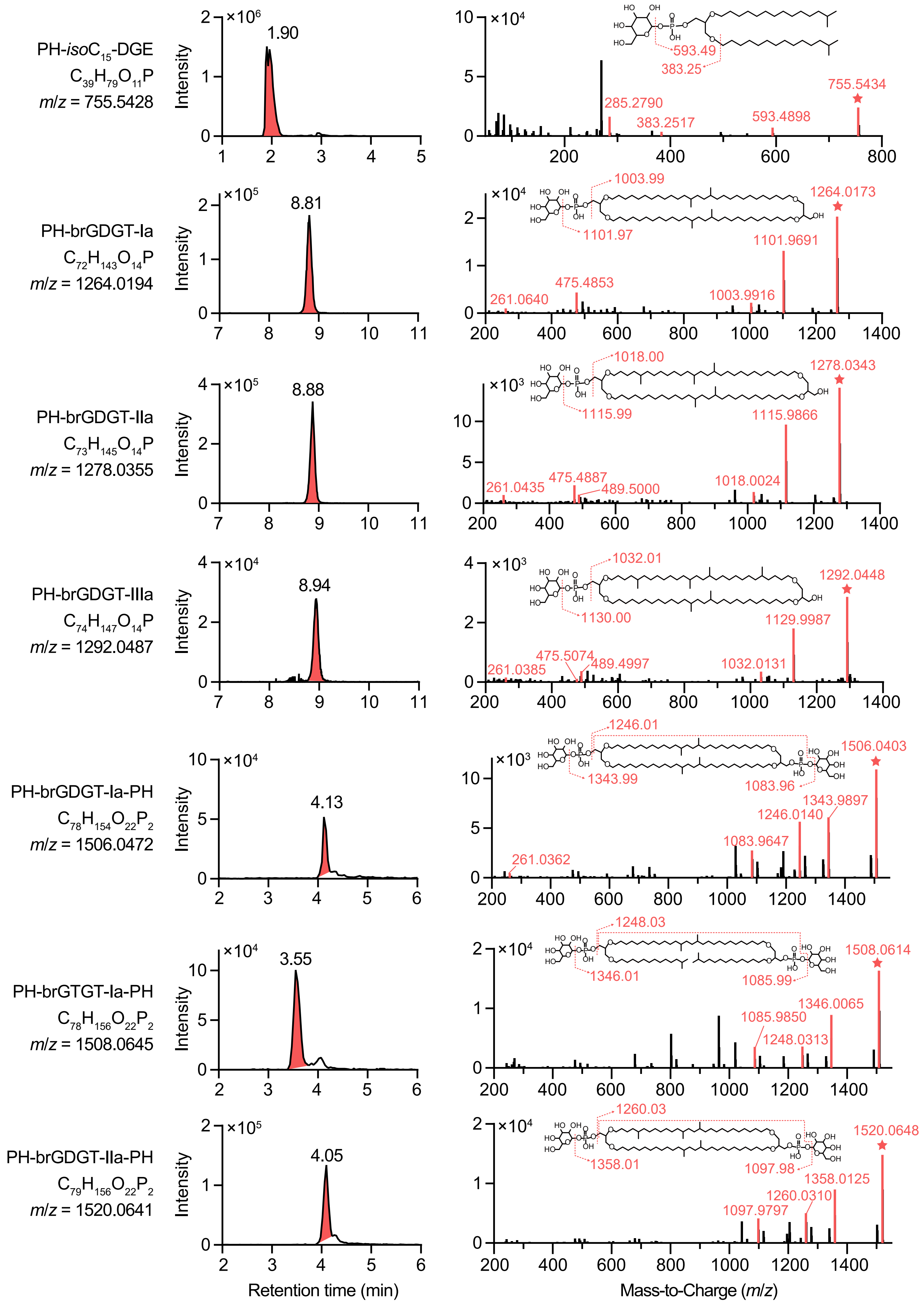


**Fig. S3.** Extracted ion chromatograms and MS² mass spectra of major IPL-brGDGT derivatives identified in strain Ellin6076 using RP–LC–HRMS. The precursor ions ($[M+H]^+$) and characteristic product ions are marked by stars and red color, respectively, and fragment positions are denoted by dash lines.

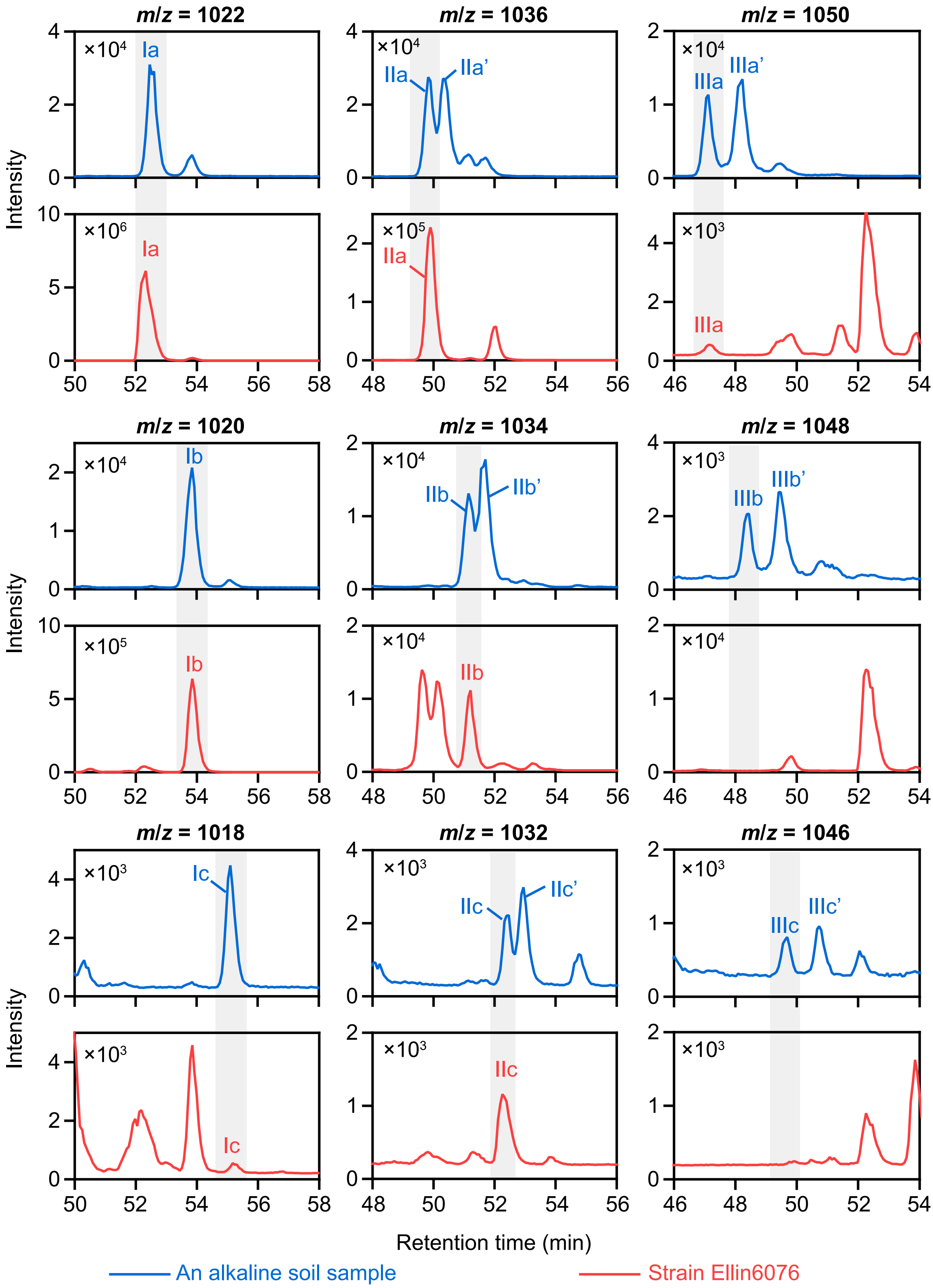


**Fig. S4.** Comparison of extracted ion chromatograms between an alkaline soil and the culture of strain Ellin6076. The strain was cultured at the optimal growth conditions (25 °C, pH 5.5). The soil sample contains both $C_5$- and $C_6$-methylated brGDGTs and was used as a reference for the determination of methyl position. The quote marks (') indicate the $C_6$-methylated isomers. Both samples were detected in the same batch using the SIM mode of NP–LC–MS. Only $C_5$-methylated brGDTGs are identified in the strain Ellin6076 according to the retention time.

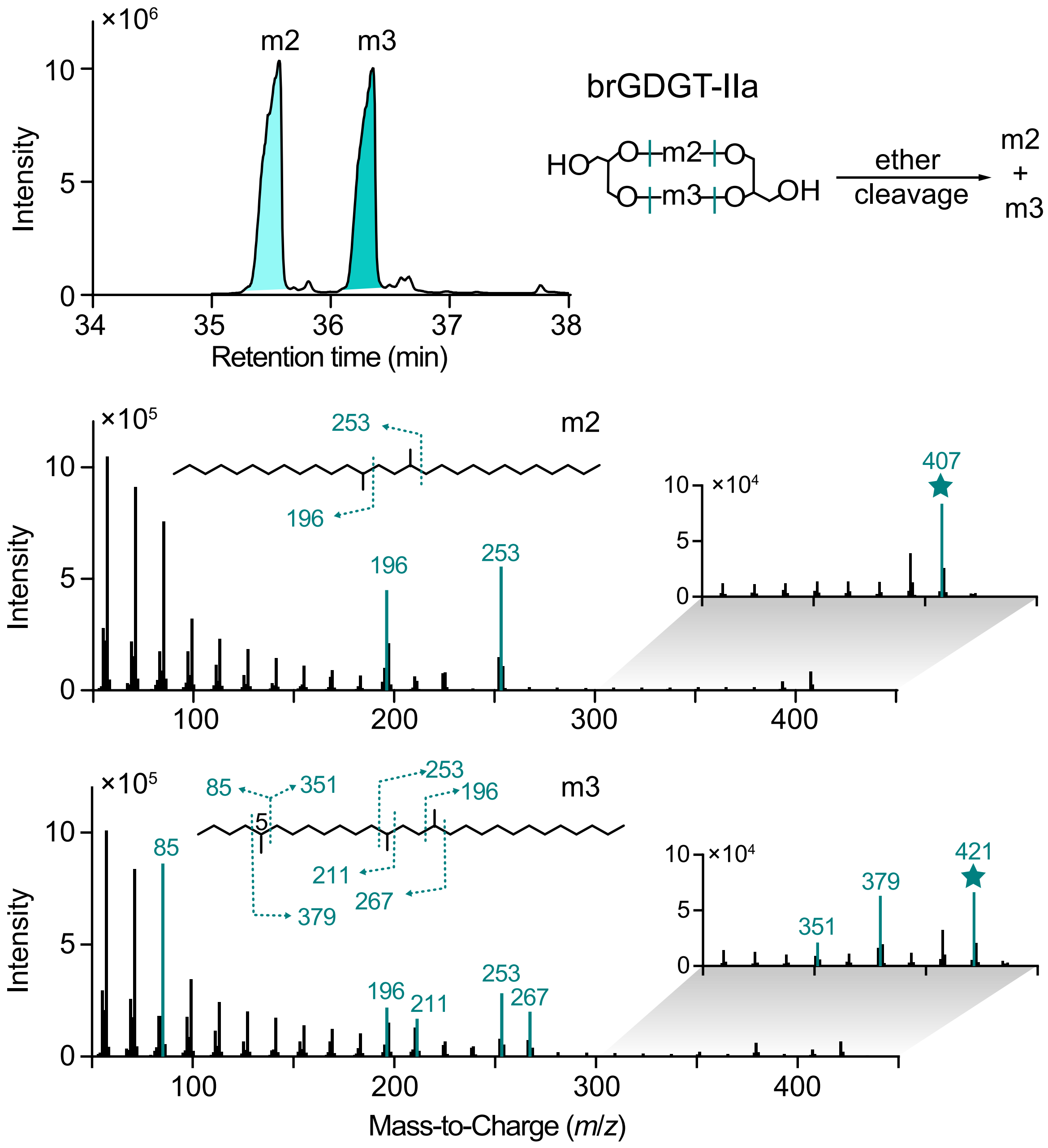


**Fig. S5.** The partial total ion chromatogram and mass spectra of brGDGT-IIa derived alkyl chains analyzed by GC–MS. The m2 and m3 refer to the alkyl chains with 2 and 3 outer methyl groups, respectively. The precursor ions ([$M^+$–15]) are marked with stars. The characteristic product ions are marked by colors and fragment positions are denoted by dash lines. The detection of m3 (5,13,16-trimethyloctacosane) confirms the $C_5$-methylation in Ellin6076 brGDGTs.

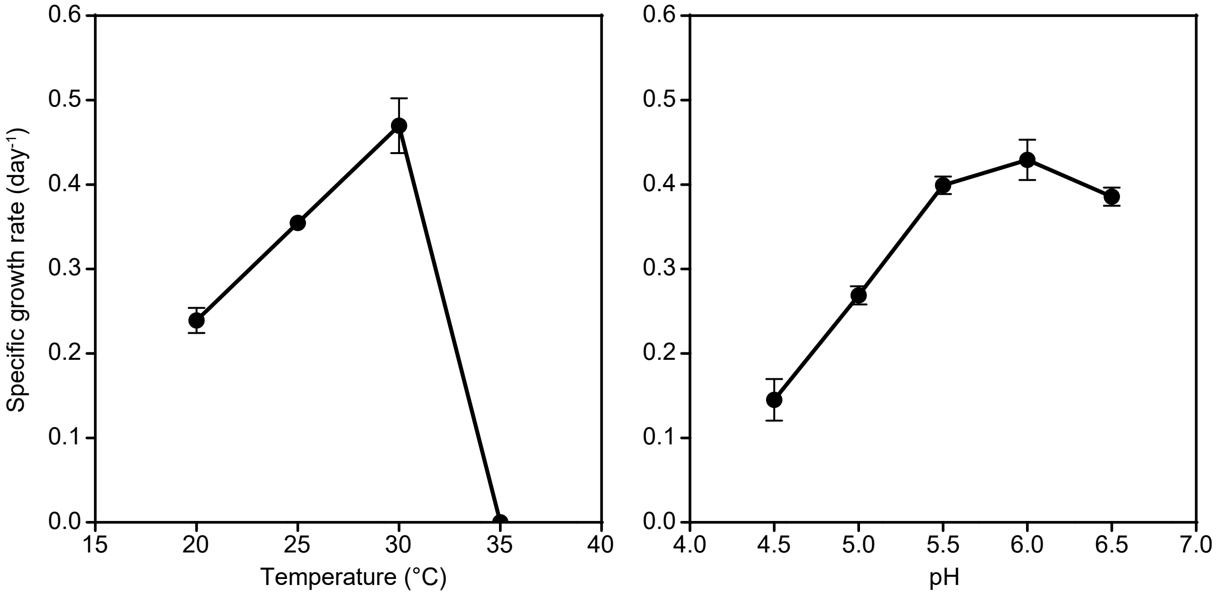


**Fig. S6.** Specific growth rate ($day^{-1}$) of strain Ellin6076 cultured under different temperature conditions (left) and pH conditions (right).

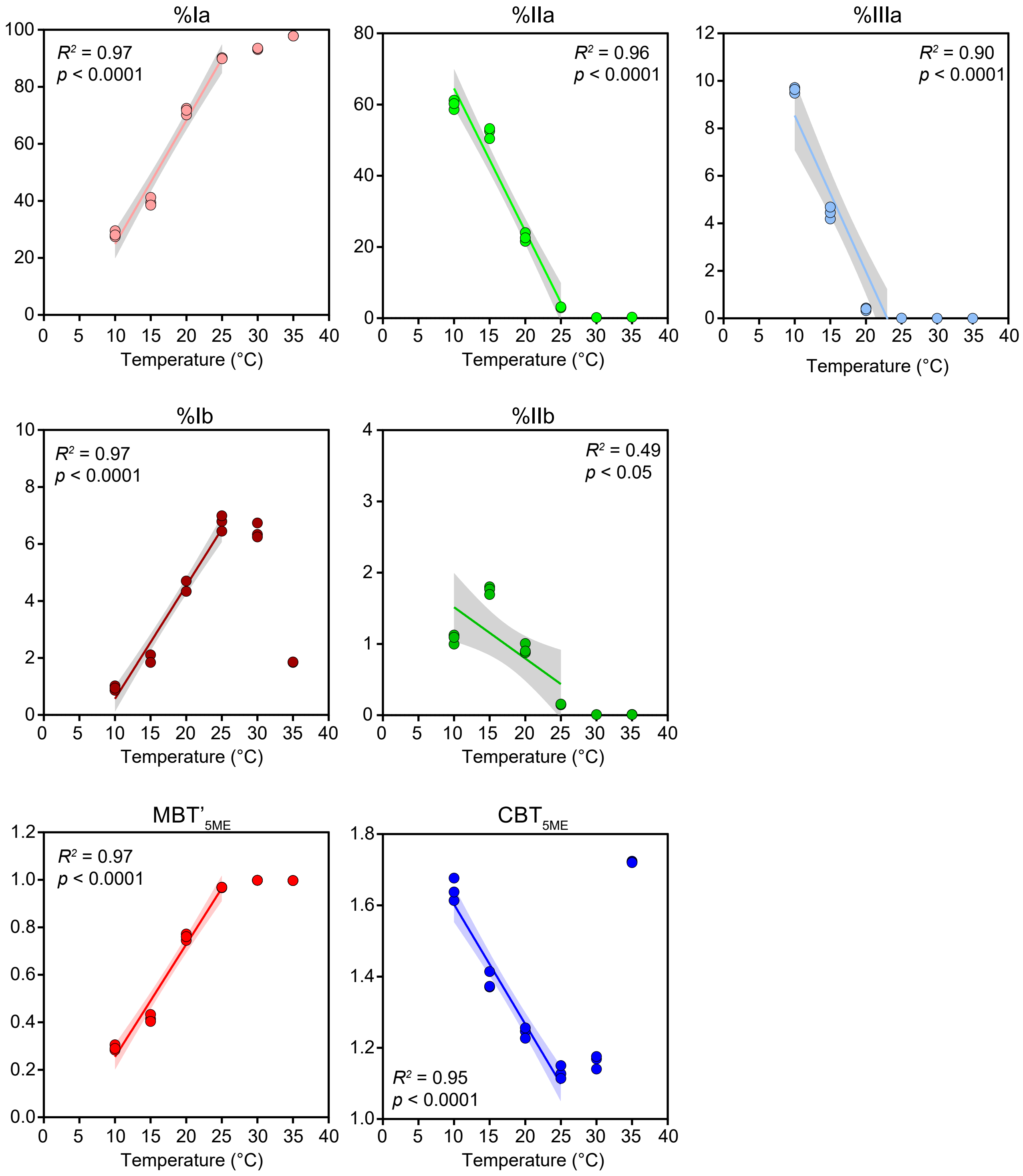


**Fig. S7.** The variation in fractional abundance (%) of major brGDGTs and brGDGT-based proxies in the cultures of strain Ellin6076 grown under different temperature conditions. The shaded area indicates 95% confidence interval and the data at 30 °C and 35 °C are excluded from the linear regression.

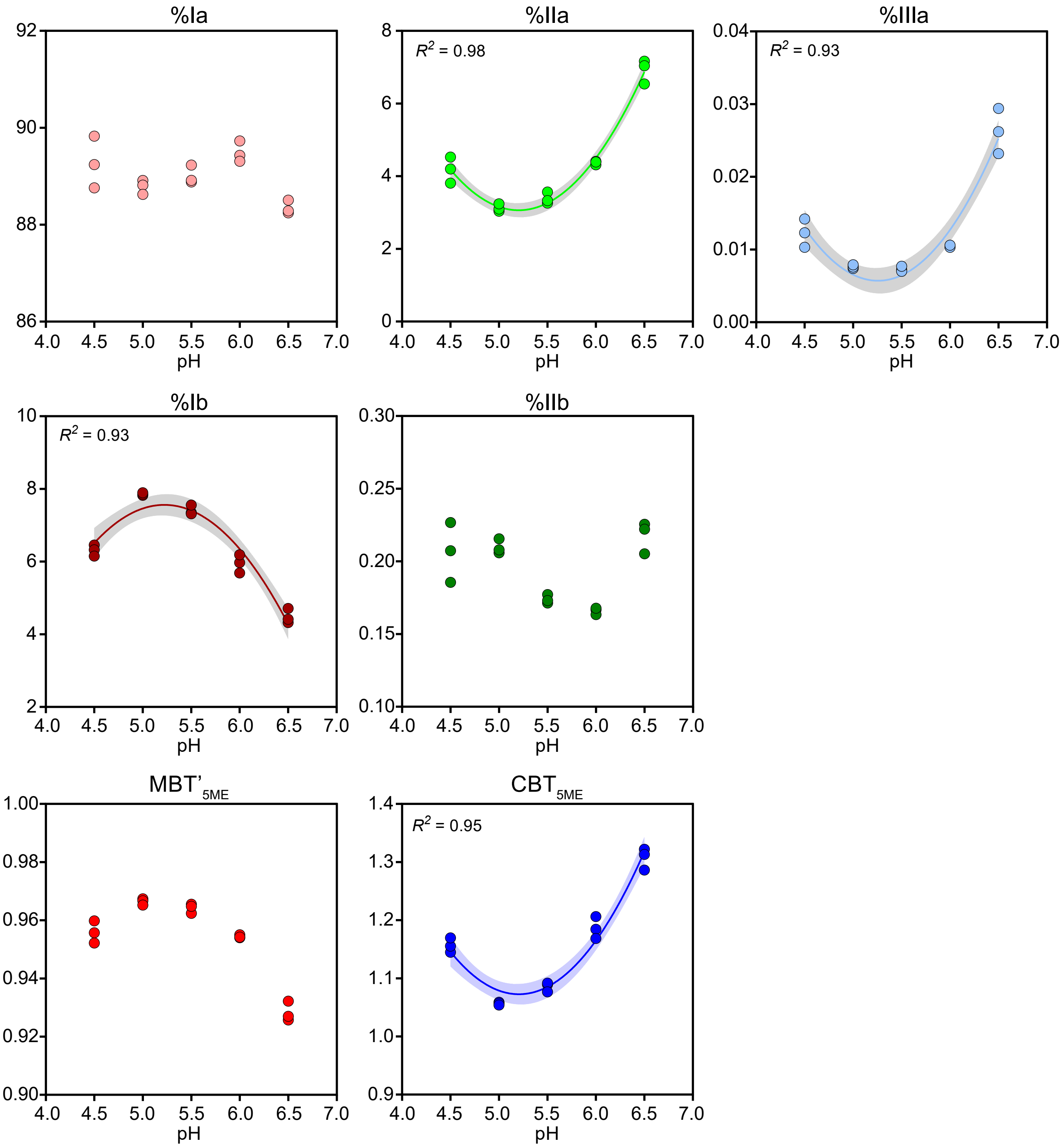


**Fig. S8.** The variation in fractional abundance (%) of major brGDGTs and brGDGT-based proxies in the cultures of strain Ellin6076 grown under different pH conditions. The shaded area indicates 95% confidence interval of second order polynomial regression.

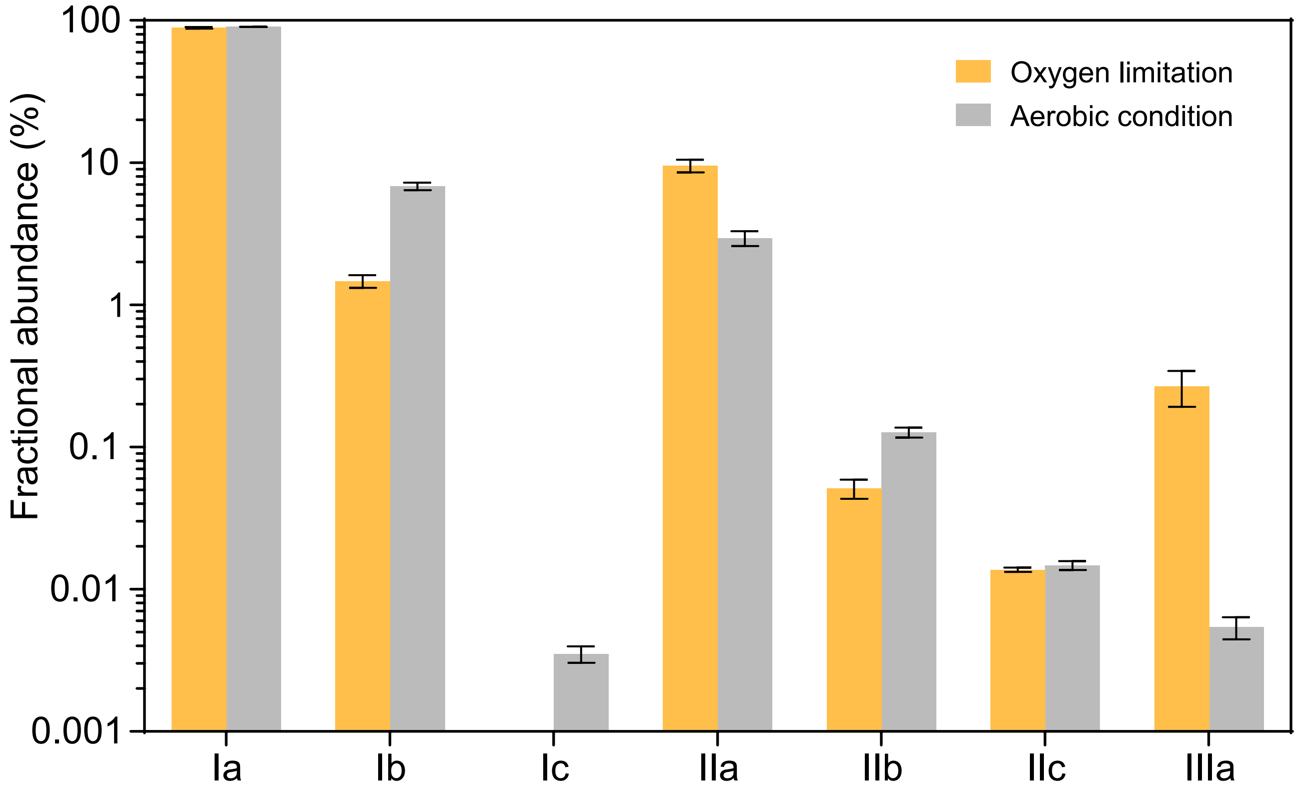


**Fig. S9.** The comparison of fractional abundance (%) of brGDGTs in the cultures of strain Ellin6076 grown under oxygen limitation and aerobic condition, respectively.

**Table S1.** The cellular contents (fg/cell) of lipid compounds in the culture of strain Ellin6076. The italics indicate the major lipid components. The percentage in the parentheses represents the fractional abundance of the lipid component in the total lipids. Each replicate sample is displayed.

| | Culture sample-1 | Culture sample-2 | Culture sample-3 |
|---|---|---|---|
| *Regular fatty acids* | | | |
| $iC_{14}$ | 0.01 | 0.01 | 0.01 |
| $nC_{14}$ | 0.01 | 0.02 | 0.01 |
| un-br$C_{15}$-1 | 1.34 | 2.41 | 1.30 |
| un-br$C_{15}$-2 | 0.07 | 0.10 | 0.07 |
| $iC_{15}$ | 3.47 | 6.25 | 3.33 |
| $aC_{15}$ | 0.01 | 0.02 | 0.01 |
| $nC_{15:1}$ | 0.01 | 0.02 | 0.01 |
| $nC_{15}$ | 0.01 | 0.01 | 0.01 |
| $iC_{16}$ | 0.03 | 0.06 | 0.02 |
| $aC_{16}$ | 0.03 | 0.05 | 0.03 |
| $nC_{16:1\text{-}1}$ | 0.90 | 1.56 | 0.83 |
| $nC_{16:1\text{-}2}$ | 0.04 | 0.04 | 0.03 |
| $nC_{16}$ | 0.19 | 0.45 | 0.14 |
| $nC_{17:1}$-1 | 3.90 | 7.03 | 3.65 |
| $nC_{17:1}$-2 | 0.28 | 0.31 | 0.20 |
| $iC_{17}$ | 0.41 | 0.66 | 0.33 |
| $aC_{17}$ | 0.04 | 0.07 | 0.04 |
| $nC_{17:1}$ | 0.08 | 0.13 | 0.07 |
| $nC_{17}$ | 0.02 | 0.03 | 0.02 |
| $nC_{18:1}$-1 | 0.06 | 0.10 | 0.05 |
| $nC_{18:1}$-2 | 0.03 | 0.04 | 0.02 |
| $nC_{18}$ | 0.08 | 0.19 | 0.07 |
| $nC_{19}$ | 0.01 | 0.02 | 0.01 |
| $nC_{20:1}$ | 0.02 | 0.04 | 0.02 |
| $nC_{20}$ | 0.07 | 0.14 | 0.08 |
| *Other lipids* | | | |
| $iC_{14}$-OH | 0.02 | 0.03 | 0.01 |
| $nC_{14}$-OH | 0.00 | 0.01 | 0.00 |
| $nC_{15}$-OH | 0.01 | 0.01 | 0.01 |
| 2-OH $C_{14}$ | 0.00 | 0.01 | 0.01 |
| 2-OH $C_{15}$ | 0.09 | 0.15 | 0.12 |
| $C_{18}$-OH | 0.02 | 0.03 | 0.01 |
| 3-OH $C_{17}$ | 0.04 | 0.10 | 0.09 |
| hop-17(21)-ene | 0.04 | 0.06 | 0.05 |
| *iso*$C_{15}$ *glycerol ethers* | | | |
| *iso*$C_{15}$-MGE | 0.27 | 0.53 | 0.36 |
| *iso*$C_{15}$-DGE | 0.67 | 1.26 | 0.61 |
| *brGDGTs* | | | |
| brGTDT-Ia | 0.08 | 0.16 | 0.10 |
| brGTDT-IIa | 0.01 | 0.02 | 0.01 |
| brGTDT-IIIa | 0.01 | 0.03 | 0.02 |
| brGDGT-Ia | 18.52 | 38.55 | 24.45 |
| brGDGT-Ib | 1.42 | 3.07 | 1.72 |
| brGDGT-Ic | 0.00 | 0.00 | 0.00 |
| brGDGT-IIa | 0.56 | 1.18 | 0.91 |
| brGDGT-IIb | 0.02 | 0.05 | 0.04 |
| brGDGT-IIc | 0.00 | 0.01 | 0.00 |
| brGDGT-IIIa | 0.00 | 0.00 | 0.00 |
| *Summed* | | | |
| *Regular fatty acids* | 11.12 (33.8%) | 19.77 (30.4%) | 10.36 (26.7%) |
| *Other lipids* | 0.23 (0.7%) | 0.40 (0.6%) | 0.30 (0.8%) |
| *iso*$C_{15}$ *glycerol ethers* | 0.93 (2.8%) | 1.79 (2.8%) | 0.97 (2.5%) |
| *brGDGTs* | 20.64 (62.7%) | 43.08 (66.2%) | 27.25 (70.1%) |

**Table S2.** The information and brGDGT data of the independent culturing experiments under different growth temperature, pH and oxygen conditions. Each replicate sample is displayed. N.D. = not detected.

| | | | | brGDGTs (fg/cell) | fractional abundance of individual brGDGTs (%) | | | | | | | Proxies | |
|---|---|---|---|---|---|---|---|---|---|---|---|---|---|
| Temperature (°C) | pH | Oxygen condition | Sample | | Ia | Ib | Ic | IIa | IIb | IIc | IIIa | $MBT'_{5ME}$ | $CBT_{5ME}$ |
| 10 | 5.5 | Aerobic | T10-1 | 37.7 | 27.44 | 0.87 | N.D. | 61.21 | 1.00 | 0.00 | 9.48 | 0.28 | 1.68 |
| 10 | 5.5 | Aerobic | T10-2 | 52.6 | 29.53 | 1.02 | N.D. | 58.61 | 1.12 | 0.00 | 9.71 | 0.31 | 1.61 |
| 10 | 5.5 | Aerobic | T10-3 | 58.0 | 28.05 | 0.94 | N.D. | 60.27 | 1.09 | 0.00 | 9.64 | 0.29 | 1.64 |
| 15 | 5.5 | Aerobic | T15-1 | 35.8 | 39.34 | 2.11 | 0.00 | 52.55 | 1.80 | 0.01 | 4.19 | 0.41 | 1.37 |
| 15 | 5.5 | Aerobic | T15-2 | 27.4 | 41.19 | 2.11 | 0.00 | 50.46 | 1.77 | 0.01 | 4.45 | 0.43 | 1.37 |
| 15 | 5.5 | Aerobic | T15-3 | 20.2 | 38.51 | 1.84 | 0.00 | 53.25 | 1.69 | 0.01 | 4.69 | 0.40 | 1.41 |
| 20 | 5.5 | Aerobic | T20-1 | 34.2 | 70.19 | 4.34 | 0.00 | 24.01 | 1.01 | 0.01 | 0.43 | 0.75 | 1.25 |
| 20 | 5.5 | Aerobic | T20-2 | 32.9 | 72.47 | 4.70 | 0.00 | 21.60 | 0.88 | 0.01 | 0.33 | 0.77 | 1.23 |
| 20 | 5.5 | Aerobic | T20-3 | 30.4 | 71.76 | 4.33 | 0.00 | 22.59 | 0.90 | 0.01 | 0.40 | 0.76 | 1.26 |
| 25 | 5.5 | Aerobic | T25-1 | 44.4 | 90.12 | 6.79 | 0.00 | 2.92 | 0.15 | 0.01 | 0.01 | 0.97 | 1.13 |
| 25 | 5.5 | Aerobic | T25-2 | 29.2 | 90.20 | 6.45 | 0.00 | 3.16 | 0.16 | 0.01 | 0.01 | 0.97 | 1.15 |
| 25 | 5.5 | Aerobic | T25-3 | 47.3 | 89.84 | 6.99 | 0.00 | 2.99 | 0.15 | 0.01 | 0.01 | 0.97 | 1.11 |
| 30 | 5.5 | Aerobic | T30-1 | 49.8 | 93.06 | 6.74 | 0.00 | 0.17 | 0.01 | 0.01 | 0.00 | 1.00 | 1.14 |
| 30 | 5.5 | Aerobic | T30-2 | 49.8 | 93.47 | 6.34 | 0.00 | 0.17 | 0.01 | 0.01 | 0.00 | 1.00 | 1.17 |
| 30 | 5.5 | Aerobic | T30-3 | 49.0 | 93.55 | 6.25 | 0.00 | 0.18 | 0.01 | 0.01 | 0.00 | 1.00 | 1.18 |
| 35 | 5.5 | Aerobic | T35-1 | 29.9 | 97.84 | 1.84 | N.D. | 0.29 | 0.01 | 0.02 | 0.00 | 1.00 | 1.72 |
| 35 | 5.5 | Aerobic | T35-2 | 23.7 | 97.84 | 1.85 | N.D. | 0.28 | 0.01 | 0.02 | 0.00 | 1.00 | 1.72 |
| 35 | 5.5 | Aerobic | T35-3 | 36.0 | 97.84 | 1.86 | N.D. | 0.28 | 0.01 | 0.01 | 0.00 | 1.00 | 1.72 |
| 25 | 4.5 | Aerobic | pH4.5-1 | 20.0 | 88.76 | 6.46 | 0.00 | 4.53 | 0.23 | 0.01 | 0.01 | 0.95 | 1.14 |
| 25 | 4.5 | Aerobic | pH4.5-2 | 17.2 | 89.24 | 6.33 | 0.00 | 4.20 | 0.21 | 0.01 | 0.01 | 0.96 | 1.16 |
| 25 | 4.5 | Aerobic | pH4.5-3 | 20.2 | 89.83 | 6.15 | 0.00 | 3.81 | 0.19 | 0.01 | 0.01 | 0.96 | 1.17 |
| 25 | 5.0 | Aerobic | pH5.0-1 | 37.6 | 88.91 | 7.82 | 0.00 | 3.04 | 0.21 | 0.01 | 0.01 | 0.97 | 1.06 |
| 25 | 5.0 | Aerobic | pH5.0-2 | 36.3 | 88.81 | 7.85 | 0.00 | 3.10 | 0.21 | 0.01 | 0.01 | 0.97 | 1.06 |
| 25 | 5.0 | Aerobic | pH5.0-3 | 31.8 | 88.63 | 7.90 | 0.00 | 3.24 | 0.22 | 0.01 | 0.01 | 0.97 | 1.05 |
| 25 | 5.5 | Aerobic | pH5.5-1 | 39.2 | 88.88 | 7.35 | 0.00 | 3.56 | 0.18 | 0.01 | 0.01 | 0.96 | 1.09 |
| 25 | 5.5 | Aerobic | pH5.5-2 | 34.3 | 89.23 | 7.32 | 0.00 | 3.26 | 0.17 | 0.01 | 0.01 | 0.97 | 1.09 |
| 25 | 5.5 | Aerobic | pH5.5-3 | 35.1 | 88.92 | 7.56 | 0.00 | 3.33 | 0.17 | 0.01 | 0.01 | 0.96 | 1.08 |
| 25 | 6.0 | Aerobic | pH6.0-1 | 34.0 | 89.43 | 5.97 | 0.00 | 4.41 | 0.17 | 0.01 | 0.01 | 0.95 | 1.18 |
| 25 | 6.0 | Aerobic | pH6.0-2 | 33.7 | 89.31 | 6.19 | 0.00 | 4.31 | 0.16 | 0.01 | 0.01 | 0.95 | 1.17 |
| 25 | 6.0 | Aerobic | pH6.0-3 | 36.9 | 89.73 | 5.69 | 0.00 | 4.39 | 0.17 | 0.01 | 0.01 | 0.95 | 1.21 |
| 25 | 6.5 | Aerobic | pH6.5-1 | 47.9 | 88.51 | 4.71 | 0.00 | 6.54 | 0.21 | 0.01 | 0.02 | 0.93 | 1.29 |
| 25 | 6.5 | Aerobic | pH6.5-2 | 52.7 | 88.24 | 4.32 | 0.00 | 7.17 | 0.23 | 0.01 | 0.03 | 0.93 | 1.32 |
| 25 | 6.5 | Aerobic | pH6.5-3 | 48.9 | 88.28 | 4.41 | 0.00 | 7.04 | 0.22 | 0.01 | 0.03 | 0.93 | 1.31 |
| 25 | 5.5 | Oxygen limitation | OL-1 | 9.6 | 89.47 | 1.34 | N.D. | 8.92 | 0.04 | 0.01 | 0.21 | 0.91 | 1.85 |
| 25 | 5.5 | Oxygen limitation | OL-2 | 4.6 | 89.30 | 1.43 | N.D. | 8.96 | 0.05 | 0.01 | 0.24 | 0.91 | 1.82 |
| 25 | 5.5 | Oxygen limitation | OL-3 | 12.4 | 87.33 | 1.63 | N.D. | 10.61 | 0.06 | 0.01 | 0.35 | 0.89 | 1.76 |
| 25 | 5.5 | Aerobic control | OC-1 | 20.5 | 90.20 | 6.93 | 0.00 | 2.73 | 0.12 | 0.01 | 0.00 | 0.97 | 1.12 |
| 25 | 5.5 | Aerobic control | OC-2 | 42.9 | 89.94 | 7.17 | 0.00 | 2.75 | 0.12 | 0.02 | 0.00 | 0.97 | 1.10 |
| 25 | 5.5 | Aerobic control | OC-3 | 27.1 | 90.15 | 6.34 | 0.00 | 3.35 | 0.14 | 0.01 | 0.01 | 0.96 | 1.16 |
